# Harnessing Neutrophils to Deliver Antibiotics Non-Invasively across the Tympanic Membrane for Otitis Media Treatment

**DOI:** 10.1101/2025.03.12.642923

**Authors:** Wenjing Tang, Xiaojing Ma, Clara M. Marlowe, Sophie S. Liu, Kwang-Won Park, Pengyu Chen, Haonian Shu, Rong Yang

## Abstract

Acute otitis media (AOM) is a leading cause of pediatric antibiotic prescriptions. Systemic antibiotics cause side effects and antibiotic resistance, whereas local delivery of antibiotics (directly to the middle ear) is hindered by an impermeable biological barrier, the tympanic membrane (TM). Here, we report on a liposome that hitchhikes on neutrophils to deliver antibiotics to the site of infection. Distinct from previous immune-cell-based therapies, we enable neutrophil hitchhiking via a topical application of hydroxylated liposomes, thus bypassing the invasive neutrophil harvesting procedures. The hydroxylated liposomes are opsonized by the complement protein fragments and subsequently internalized by native neutrophils. A simple topical application completely cures AOM in an established chinchilla model. It points to a low-cost and non-invasive treatment for this prevalent disease, poised to reduce pediatric antibiotic usage.

## INTRODUCTION

Acute otitis media (AOM) is the primary reason for pediatric antibiotic usage, accounting for 24% of the antibiotic prescriptions written to U.S. children (*1*). An estimated 19.5 million AOM cases in the U.S. each year lead to an annual cost of $3-4 billion from outpatient visits, emergency department/urgent care visits, and hospitalization (*2*). Globally, ∼ 80% of all children experience at least one episode of AOM by school age (*3*), making AOM one of the most common reasons for pediatrician visits around the world.

AOM is predominantly caused by bacterial infections (*4, 5*), among which nontypeable *Haemophilus influenzae* (NTHi) accounts for 30-52% of all AOM episodes globally (*6, 7*). The current mainstay treatment for AOM comprises oral antibiotics for 5-10 days (*8*). However, oral antibiotics cause acute side effects in children, including diarrhea and vomiting (*9*), and are correlated with long-term health issues like asthma and obesity (*10*). As a result, the medication is discontinued prematurely in ∼ 1/3 of U.S. children who are 6-36 months of age, frequently upon resolution of symptoms (*11*). Poor patient compliance, in turn, speeds up the development of antibiotic resistance. A treatment strategy to minimize antibiotic usage during AOM treatment could mitigate the side effects and development of drug resistance.

Here, we report on a fresh concept to enable neutrophil-based drug delivery via topical application without the need for immune cell harvesting, thereby targeting the delivery of antibiotics to the infected middle ear. Existing immune-cell-mediated therapies are incompatible with topical application. They often rely on the invasive and costly procedure of harvesting immune cells ex vivo, loading drugs, and re-introducing the immune cells back into systemic circulation (*12–14*). In contrast, we avoid invasive procedures and potential systemic exposure by discovering that neutrophils can readily penetrate the impermeable tympanic membrane (TM), providing a pathway between the outer ear canal and the infected middle ear. Our design thus harnesses neutrophils as endogenous micromotors to traffic the antibiotic cargo that is placed in the patient’s outer ear canal across the TM, targeting the site of infection (Scheme 1).

The TM, while only ∼ 100-μm thick in humans (*15*), has been a longstanding bottleneck in transtympanic drug delivery, as it is largely impermeable to most molecules, including antibiotics. Our observation of neutrophils (∼ 12 μm in diameter) throughout the TM (Fig. S1) is unprecedented despite their reported presence in skin during psoriasis and dermatophytosis (*16, 17*). Building upon this observation, we design a liposome-based drug delivery system that achieves a transtympanic antibiotic delivery efficiency of 14.4% (4∼8-fold the best-in-class formulations, which use chemical permeation enhancers to disrupt the TM (*18–20*)). The formulation, when placed in the outer ear canal of chinchillas infected by NTHi, completely eradicates AOM in all treated animals.

**Scheme 1.**
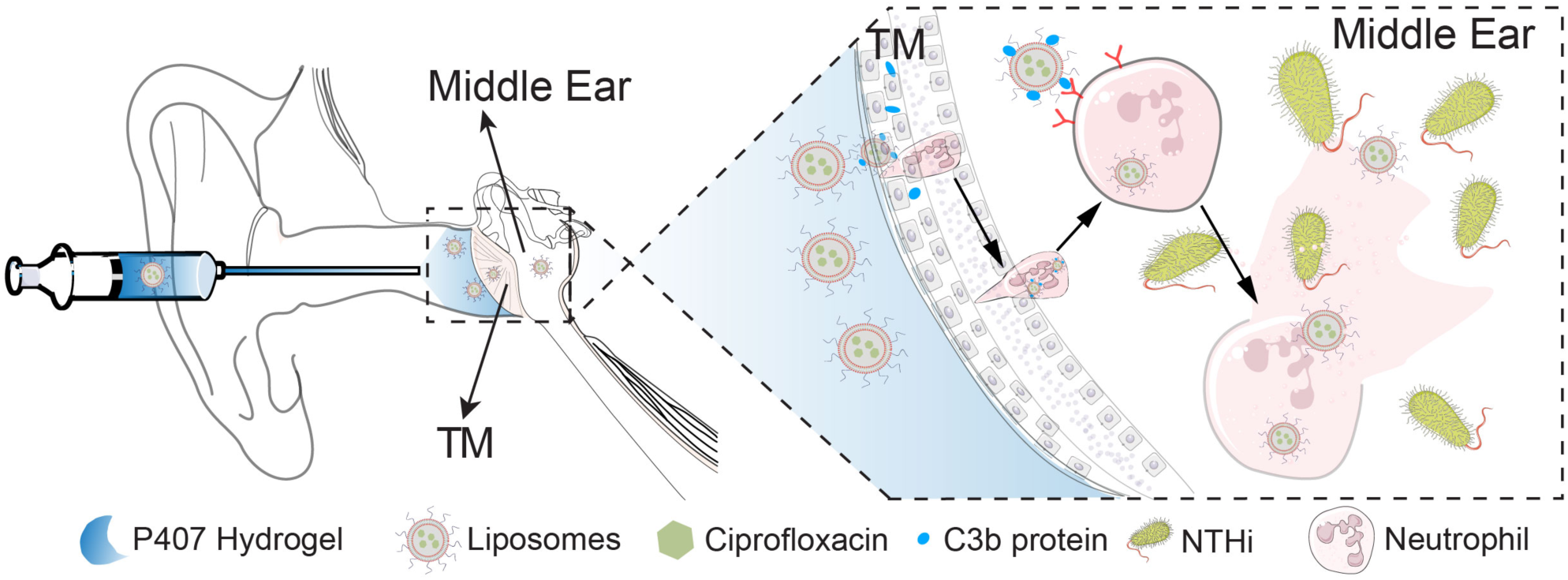
Schematic illustration of the transtympanic delivery system, leveraging the native immune response triggered by an acute otitis media (AOM) episode. The neutrophils recruited by the infection internalize the C3b-marked liposomes and traffic them across an intact tympanic membrane (TM).

The liposomes hitchhike on neutrophils by leveraging the complement cascade, a key component of the innate immunity and host defense system (*21*). During AOM, neutrophils, the most abundant leukocytes in the bloodstream (*22*), undergo chemotaxis, migrating from the bloodstream toward the middle ear (*23*). Along the way, neutrophils internalize foreign substances that have been opsonized by the complement molecules (*24*), most notably by the cleavage products of the complement 3 (C3) protein, i.e., C3b and its subsequent cleavage molecule, iC3b (*25, 26*). It has been demonstrated that opsonization could be enabled by a reaction between a short-lived thioester moiety in C3b and hydroxyl (OH) groups on foreign substances (*21, 27*). As such, we design a delivery system containing OH-bearing liposomes made of 1,2-distearoyl-sn-glycero-3-phosphoglycerol (DSPG) (Fig. 1A). The C3b fragments covalently linked to the liposome serve as biomarkers for enhanced internalization and trafficking by innate neutrophils to the infected middle ear.

**Fig. 1.**
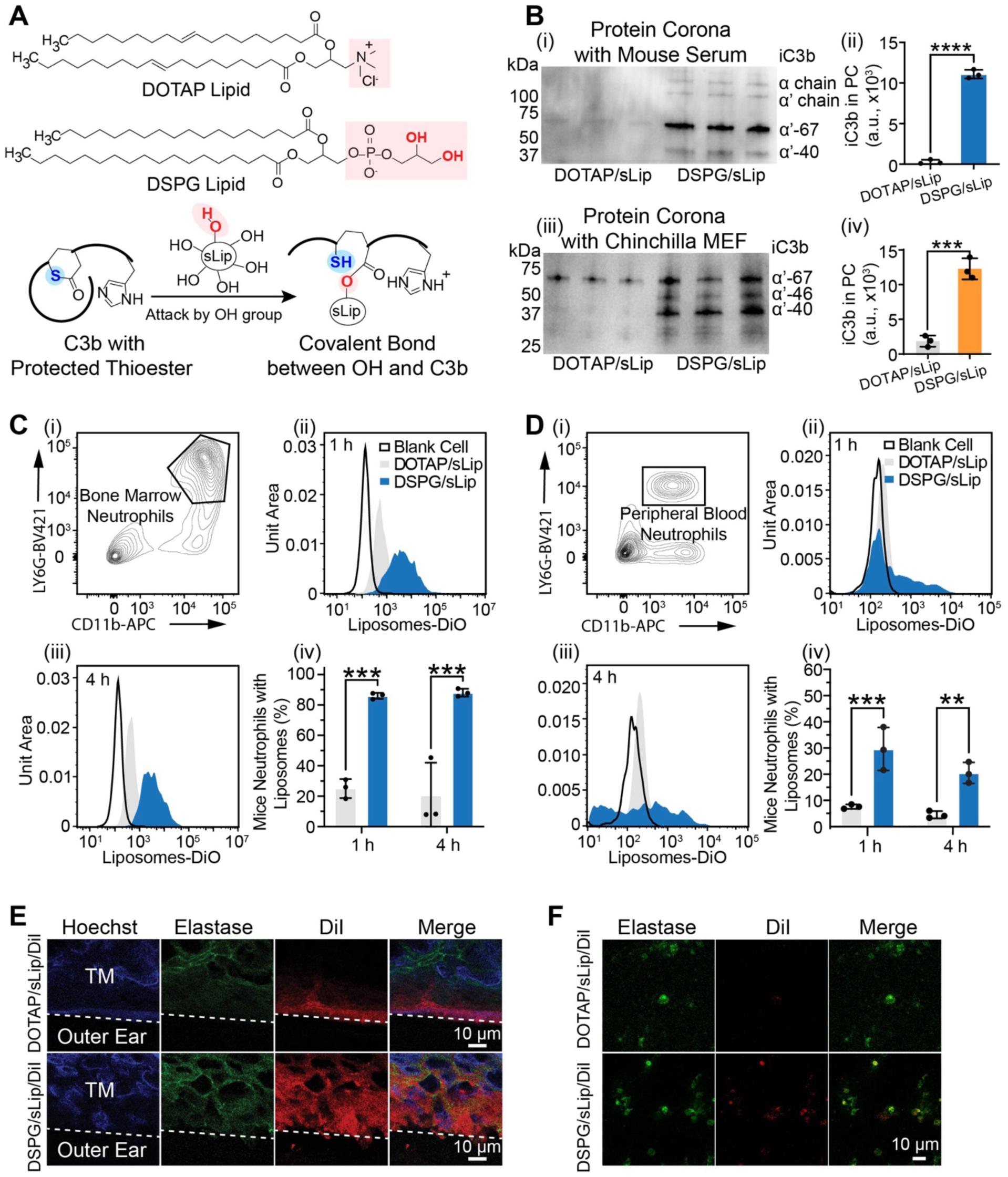
Hydroxyl-bearing liposomes traverse the tympanic membrane by hitchhiking on neutrophils, enabled via a reaction with the complement protein fragment, C3b. **(A)** Molecular structures of a non-hydroxyl-bearing lipid, i.e., dioleoyl-3-trimethylammonium propane (DOTAP), and a hydroxyl-bearing lipid, i.e., 1,2-distearoyl-sn-glycero-3-phosphoglycerol (DSPG), and the reaction between the hydroxyl groups displayed by the DSPG liposomes (DSPG/sLip) and C3b that marks DSPG/sLip for subsequent neutrophil internalization. **(B)** Western blot of the liposomes’ protein corona after incubating with BALB/c mice serum (i-ii) (\*\*\*\**P* < 0.0001) and the middle ear fluid (MEF) collected from chinchillas with AOM (iii-iv) (*P* = 0.0005) respectively, and the quantity of iC3b fragments in the protein corona of DOTAP/sLip and DSPG/sLip, respectively. **(C)** ex vivo uptake of the opsonized liposomes by the innate neutrophils harvested from the bone marrow of BALB/c mice; the identity of the neutrophils was confirmed using flow cytometry, where neutrophils were gated at CD11b+/LY6G+ (i); the liposomes were fluorescently labeled using 3,3’-Dioctadecyloxacarbocyanine (DiO), and incubated with the neutrophils ex vivo for 1 h (ii) and 4 h (iii), respectively, with the fraction of liposome-containing neutrophils quantified (iv); \*\*\**P* = 0.0003 and \*\*\**P* = 0.0002. **(D)** in vivo uptake after intravenous injection (50 mg lipid per kg body weight) of the two types of liposomes by BALB/c mice peripheral neutrophils (i) at 1h (ii) and 4 h (iii), respectively, with the fraction of liposome-containing neutrophils quantified (iv); \*\*\**P* = 0.0008 and \*\**P* = 0.0058. **(E)** Fluorescent images of the auditory bullae of chinchillas with AOM treated with the two liposomal formulations and the amounts of each liposome in the TM (\*\**P* = 0.0047), estimated from their fluorescent intensities; liposomes are fluorescently labeled using tetramethylindocarbocyanine perchlorate (DiI, red); DSPG/sLip/DiI readily traverse the TM (blue, with Hoechst staining for cell nucleus) by hitchhiking on the native neutrophils (green, with elastase staining), whereas DOTAP/sLip/DiI remain largely in the stratum corneum layer. **(F)** Fluorescent images of the MEF of chinchillas with AOM treated with the two liposomal formulations and the amounts of each liposome in the MEF (\**P* = 0.0454), estimated from their fluorescent intensities. n = 3 for each group; data are presented as mean ± s.d.; scale bar, 10 µm; *P* values were determined by two-tailed unpaired Student’s t-tests; \**P* < 0.05; \*\**P* < 0.01; \*\*\**P* < 0.001 and **** *P* < 0.0001. See Fig. S4 for the gating strategy of neutrophil subsets in (C) and (D); See Fig. S5 for the estimated fluorescence intensity of liposomes on TM (E) and in the neutrophils from MEF (F).

To treat AOM, we chose ciprofloxacin as the antibiotic because it is commonly used in the current treatments (e.g., applied to the middle ear via a perforated TM) (*28, 29*). The ciprofloxacin-encapsulating DSPG liposomes were loaded into a thermoresponsive hydrogel made of Poloxamer 407 (P407). Importantly, the formulation is a solution under room temperature for ease of administration into the outer ear canal and gels firmly upon reaching the TM. As such, a single dose (e.g., applied in the pediatrician’s office) could sustain the release of antibiotics over the 7-day treatment, avoiding the premature discontinuation of the treatment. The neutrophil-based delivery enables the eradication of AOM using a small amount of antibiotics, i.e., 0.5 mg ciprofloxacin, corresponding to 6.25% of that in the best-in-class transtympanic formulation (*19*) and 0.5% of that used in systemic AOM treatment (*30*). The local administration and high transtympanic delivery efficiency combined minimize antibiotic exposure during AOM treatment.

The strategy of leveraging the complement system to harness innate neutrophils as a drug delivery vehicle is unprecedented. To illustrate the critical role played by the complement system, we treated the infected chinchillas with cobra venom factor (CVF), which has been shown to deplete complement proteins (*31, 32*), prior to receiving the liposomes. Merely 25% of the animals cleared their AOM, in contrast to the 100% cure rate in the ones not treated with CVF. This strategy enables targeted delivery without the requirement for specific targeting biomolecules, such as antibodies, that are costly to deploy for such a prevalent disease. The transtympanic delivery platform has the potential to reduce pediatric antibiotic usage globally, thus mitigating acute and long-term health issues related to antibiotic exposure early in life.

## RESULTS

### Hydroxyl-bearing liposomes traverse the TM of infected animals by hitchhiking on innate neutrophils via the opsonization by complement molecules

We performed the ex vivo and in vivo experiments using a chinchilla model of AOM, which was induced by direct inoculation of NTHi into the auditory bullae. *Chinchilla Lanigera* was used as it is a preclinical model widely accepted for the pathophysiology and treatment of AOM (*33, 34*). The liposome design is based on our observation of neutrophils throughout the TM during an active episode of AOM. While inflammation or neutrophils are absent from the TM under normal homeostasis (Pristine TM in Fig. 6A), a considerable number of neutrophils were observed in the TM and auditory bullae (Fig. S1, A and C), excised at 48 hours or 72 hours post-infection (to be consistent with the ex vivo and in vivo efficacy tests described in later sections). There are ∼ 30 neutrophils visible in the histology image shown, permeating the entire TM structure. Neutrophils were also captured in the middle ear fluid (MEF) (Fig. S1, B and D), which is consistent with prior reports of a neutrophil level of 40-300 cells in the middle ear cavity following an NTHi inoculation (*35, 36*). Based on a quantitative analysis of histological images from a 280 µm × 370 µm area, the neutrophil levels at 48 h and 72 h bear no significant difference (Fig. S1E). The identification of neutrophils in the TM during AOM informed the design of the liposomal formulation, as detailed below.

We hypothesize that liposomes bearing hydroxyl moieties experience enhanced opsonization by C3b fragments, which marks the liposome for internalization as a way to “load” the liposomes into neutrophils for transtympanic delivery. To verify the presence of the C3b fragments in animals with AOM, we analyzed their serum and MEF using western blot (Fig. S2, B to C). Note that an anti-guinea pig complement antibody was used in the western blot due to an incomplete protein database on *Chinchilla Lanigera*; similar methods have been used in the literature (*37–39*). The western blot identified the C3α (120 kDa) and iC3b α’ fragments (67 kDa, 46 kDa, and/or 40 kDa) in both fluids. The molecular weight of the C3 fragments was consistent with our prediction based on the known C3 cleavage mechanism via the alternative activation pathway (Fig. S2A) (*37, 40*).

To assess the effectiveness of the hitchhiking, we compared hydroxyl-bearing liposomes made of DSPG and non-hydroxyl-bearing ones made of dioleoyl-3-trimethylammonium propane (DOTAP) (Fig. 1A). DOTAP was chosen because it is positively charged and thus considered effective for penetrating the stratum corneum (the most impermeable structure in the TM) (*41, 42*), which is negatively charged (*43*). The liposomes were made in a stealth fashion, denoted as sLip, by incorporating polyethylene glycol (PEG) (at 5% molar ratio). Dynamic light scattering (DLS) measured an average diameter of 127.4 ± 5.2 nm and an average surface charge of –20.1 ± 0.4 mv for the DSPG liposomes (DSPG/sLip), and 134.2 ± 1.5 nm and +26.8 ± 1.5 mv, respectively, for the DOTAP liposomes (DOTAP/sLip) (Fig. S3 and Table S1).

The enhanced opsonization of the hydroxyl-bearing liposomes was illustrated by analyzing their protein corona using western blot (*44–47*). The protein corona was formed by incubating the liposomes with MEF harvested from infected animals. As shown in Fig. 1B-iii to iv, C3b fragments strongly bind to DSPG/sLip but not to DOTAP/sLip. The amount of the complement biomarkers displayed on the surface of DSPG/sLip is 6.6-fold that on DOTAP/sLip (*P* = 0.0005). Similar observations were made when the liposomes were incubated with the serum of chinchillas with AOM (Fig. S2, D to E). The C3b fragments are enriched in the protein corona of DSPG/sLip (*P* = 0.0082 v.s. DOTAP/sLip), as the complement cascade is a systemic reaction. To provide direct evidence that the hydroxylated liposomes (DSPG/sLip) are internalized by innate neutrophils, we adopted a BALB/c mouse model and assessed the neutrophil uptake of liposomes. We chose this mouse model because it is well-established for neutrophil harvesting and characterization (*48, 49*). These experiments cannot be performed in chinchillas because many key antibodies that are required to label neutrophils are currently missing for chinchillas. Opsonization of the hydroxylated liposomes by the complement cascade in BALB/c mice was illustrated by incubating the two types of liposomes with mouse serum for 1 hour at 37°C, followed by western blot to analyze their protein corona (Fig. 1B-i to ii). On the hydroxylated liposomes, i.e., DSPG/sLip, several iC3b fragments were identified, i.e., α’-67 and α’-40 (whereas α’-46 is an intermediate fragment that is not consistently captured; please refer to Fig. S2A for the detailed complement protein cleavage process). In contrast, the protein corona of DOTAP/sLip did not contain those complement protein fragments. The amount of the C3b fragments displayed on the surface of DSPG/sLip is 40-fold that on DOTAP/sLip (\*\*\*\**P* < 0.0001).

To illustrate neutrophils’ internalization of liposomes, we isolated fresh neutrophils from the BALB/c mouse bone marrow and incubated them with opsonized liposomes, labeled with 3,3’-dioctadecyloxacarbocyanine (DiO), for 1 hour and 4 hours, respectively. Using flow cytometry, the harvested neutrophils were gated at CD11b+/LY6G+ to confirm the cell type (*50*) (Fig. 1C-i and Fig. S4A), and the fraction of neutrophils (CD11b+/LY6G+) that contain liposomes (DiO) was quantified for each type of liposome (Fig. 1C-ii to iv). Neutrophils incubated with PBS (no liposome) were shown in Fig. 1C-ii to iii as a control group, which showed DiO signals below the threshold (i.e., 10^3^). At the end of the 1-hour incubation, 86.0 ± 2.0% neutrophils internalized DSPG/sLip (Fig. 1C-iv), as shown by the fraction of the incubated neutrophils that exhibited DiO signal above 10^3^ (Fig. 1C-ii); whereas for DOTAP/sLip, merely 25.0 ± 6.3% of the incubated neutrophils internalized the liposomes. Those internalization fractions remained similar after 4 hours of incubation, reaching 88.1 ± 2.6% for DSPG/sLip and 20.5 ± 21.5% for DOTAP/sLip, respectively. This result indicates that DSPG/sLip/DiO led to distinctly high internalization by neutrophils both at 1 hour (*P* = 0.0003) and 4 hours (*P* = 0.0002), compared to DOTAP/sLip/DiO.

To assess the in vivo neutrophil internalization efficiency, mice were injected intravenously with the two types of liposomes (DiO labeled), and flow cytometry was performed on peripheral blood (with erythrocytes removed) at 1 hour and 4 hours post-injection (Fig. 1D). Again, we confirmed the cell type(*50*) (i.e., CD11b+/LY6G+) (Fig. 1D-i and Fig. S4B), and calculated the fraction of neutrophils (CD11b+/LY6G+) that contain liposomes (DiO) for each type of liposomes (Fig. 1D-ii to iv). Neutrophils harvested from the peripheral blood of mice that received saline injection (without liposomes) were shown in Fig. 1D-ii to iii as a control group, in which the DiO signal remained below the threshold (i.e., 10^3^). At 1 hour post-injection, 29.6 ± 8.2% neutrophils internalized DSPG/sLip (Fig. 1D-ii to iv). Note this fraction is lower than that in the ex vivo experiment (Fig. 1C) because the opsonized liposomes can be eliminated by a variety of immune mechanisms (*51*), thus diluting its interaction with neutrophils specifically. For DOTAP/sLip, 7.6 ± 0.9% of the sampled neutrophils internalized the liposomes. These fractions decreased slightly at 4 hours post-injection, due to the further clearance of the injected liposomes by the immune system. As such, DSPG/sLip/DiO enhanced the in vivo neutrophil internalization to 4-fold that of DOTAP/sLip/DiO at 1 h post-injection and 5-fold at 4 h post-injection.

Finally, to visualize the permeation of the liposome-bearing neutrophils in the TM, liposomes were labeled with tetramethylindocarbocyanine perchlorate (DiI (red), a hydrophobic compound that self-assembles into the bilayer membrane of liposomes) and applied to the intact TMs of chinchillas with AOM (harvested at 72 hours post-infection). Immunocytochemistry analysis revealed that DSPG/sLip/DiI (red) overlap with neutrophils (with elastase staining for the antigen on the neutrophil surface (green)) throughout the TM (with Hoechst staining for the nucleus of live cells (blue)) (Fig. 1E). The amount of DSPG/sLip/DiI in the TM (indicated by the average DiI intensity) was 4.1-fold that of DOTAP/sLip/DiI (P = 0.0047) (Fig. S5A). To further illustrate that the liposome-bearing neutrophils can penetrate the entirety of the TM, a low-magnification fluorescent micrograph is provided in Fig. S6, in which the TM, as well as the outer and middle ear space, are visible. The overlapping green and red fluorescence throughout the TM hints at the hitchhiking of liposomes on neutrophils. In contrast, DOTAP/sLip/DiI predominantly accumulate in the external auditory canal (Fig. 1, E to F). DSPG/sLip/DiI were also identified in the MEF, indicating that liposome-bearing neutrophils are present in the auditory bullae (Fig. 1F). The DiI intensity of DSPG/sLip/DiI was 4.1-fold that of DOTAP/sLip/DiI (*P* = 0.0454) in the MEF (Fig. S5B).

### The hydroxyl-bearing liposomes enhance antibiotic delivery across infected TM ex vivo

The effectiveness of the transtympanic drug delivery strategy was demonstrated by quantifying the drug flux across intact TM in health or disease using excised auditory bullae, as shown in Fig. 2A. The intactness of excised TM was confirmed using electrical impedance measurements following a widely adopted procedure (*19, 52–54*). TMs with microperforations were excluded from the reported results. Each test formulation contains 0.5 mg ciprofloxacin (Cip) (at a concentration of 2.5 mg/ml).

**Fig. 2.**
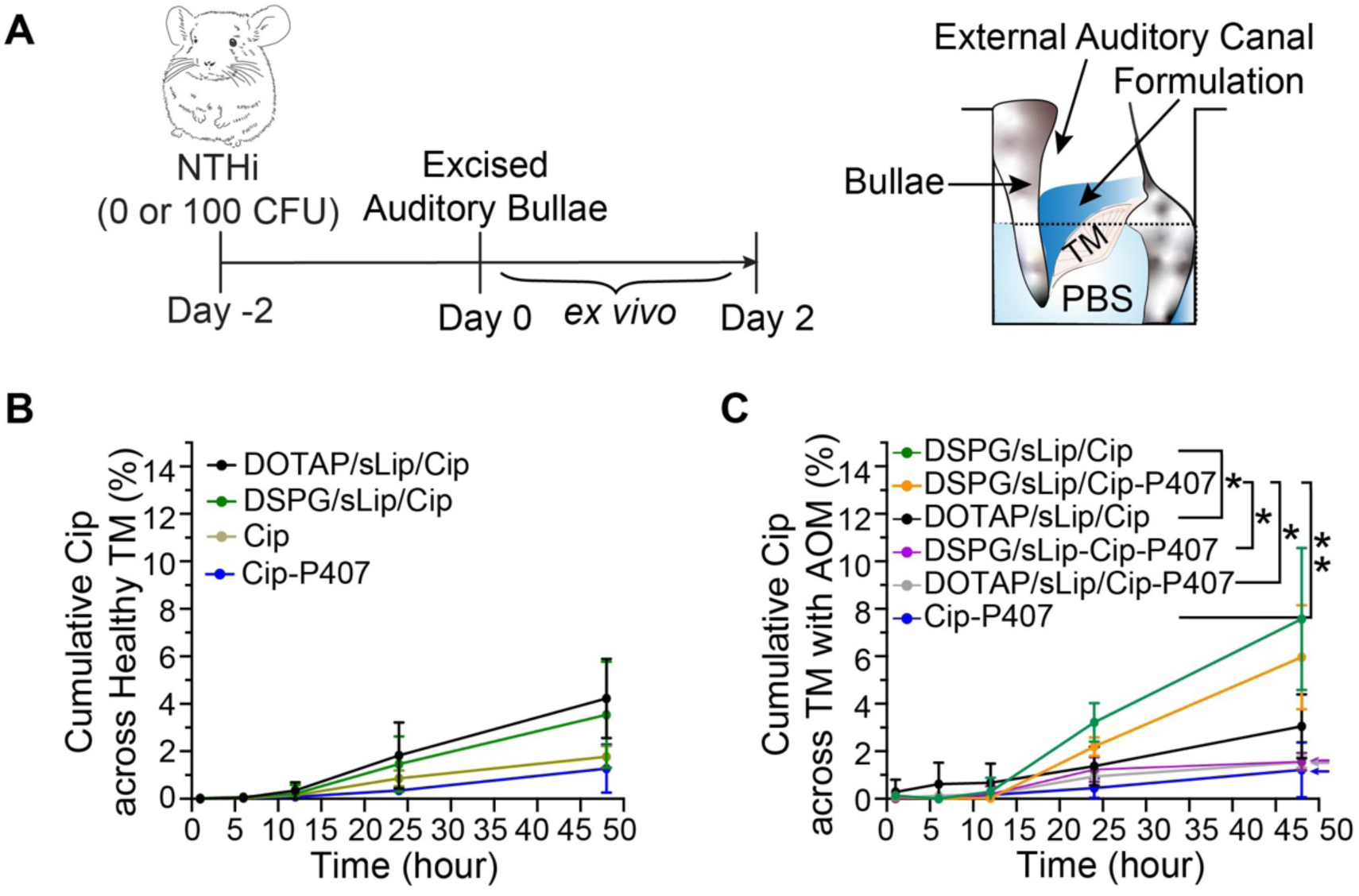
Cumulative ex vivo transfer of ciprofloxacin across the TM into a receiving chamber. (**A**) Diagram of the ex vivo protocol for assessing ciprofloxacin (Cip) permeation across the TM. A colony-forming unit (CFU) value of 0 indicates healthy animals. (**B**) Permeation (%) of Cip across the TMs of healthy chinchillas over 48 hours (n = 3). (**C**) Permeation (%) of Cip across the TMs of chinchillas with AOM over 48 hours (n = 4). DSPG/sLip-Cip-P407 (purple line) indicates a mixture of DSPG/sLip, free Cip, and P407; \**P* = 0.0146, \**P* = 0.0177, \**P* = 0.0170 and \*\**P* = 0.0097 (from left to right), calculated using one-way analysis of variance (ANOVA) with Turkey’s test; \**P* < 0.05; \*\**P* < 0.01. All formulations contain 0.5 mg Cip; in all formulations containing P407, its concentration is 15% (w/v).

Among the test formulations, DSPG/sLip/Cip shows the greatest permeability in an infected TM (Fig. 2C). Over 48 hours, 7.6 ± 3.0% of the Cip contained in this formulation permeated across TM cumulatively, representing 2.5-fold that delivered by DOTAP/sLip/Cip (3.1 ± 1.4%; P = 0.0146; Fig. S7C). This comparison contrasts with their transtympanic delivery efficiency in healthy ears (Fig. 2B and Fig. S7C), where DOTAP/sLip/Cip transported 4.2 ± 1.7% of the Cip contained in the formulation and DSPG/sLip/Cip 3.5 ± 2.2%. Data are presented as mean ± s.d. See Fig. S7C for a comparison of the cumulative Cip (%) across healthy TM and TM with OM at 48 hours.

### Drug-encapsulating liposomes are loaded into and released from a hydrogel intact without drug leakage

To cure AOM, it is critical that the antibiotics are delivered into the auditory bullae continuously over the course of 5-7 days ^42,43^. To avoid premature termination of the treatment, we designed the formulation to deliver the entire course of the regimen in a sustained fashion from a single-dose application. The sustained and local drug delivery is often achieved by loading therapeutics into a hydrogel.

We chose Poloxamer 407 (P407) because, in addition to enabling sustained drug release and ensuring biocompatibility, P407 hydrogel also undergoes reverse thermal gelation at near body temperature. P407 is a triblock polymer with poly (ethylene oxide) (PEO) endblocks and a poly(propylene oxide) (PPO) midblock. Increasing temperature drives the formation of spherical micelles composed of PPO cores and PEO shells (*57*). When a critical volume fraction of micelles is achieved, the micelles order into cubic packings, causing the low-viscosity solutions to gel (*58*). The hydrogel formulation thus remains a liquid during administration and gels upon contacting the warm TM (Scheme. 1). The ease of administration has proven key in treating young patients who are most prone to AOM.

The P407 concentration of 15% (w/v) was used in all hydrogel formulations based on an optimization of the polymer concentration. Briefly, a 10% (w/v) P407 solution did not gel at body temperature (Fig. S8A), whereas a 20% (w/v) solution had a sol-gel transition lower than 24°C (Fig. S8B), causing premature gelation and poor contact with the TM. The storage (G’) and loss (G’’) moduli of P407[15%] were below 100 Pa at room temperature (measured by linear oscillatory shear rheology at 100 rad^-1^, 1% strain, and 1°C min^-1^), which increased to 994.5 ± 370.3 Pa and 167.1 ± 23.9 Pa, respectively, at body temperature. The gelation temperature was 27-28°C.

The Cip-encapsulating liposomes were loaded into a P407 hydrogel by simple mixing (*59*). While amphiphiles are known to disrupt the gelation of P407 (*60, 61*), the addition of liposomes did not significantly alter the rheological behavior of P407[15%] (Fig. 3B). We attributed the unchanged gelation to the presence of PEG chains distributed across the surface of the liposomes (5% molar ratio in the liposomes, which has been estimated to be equivalent to 50 Å of thickness) (*62*). The storage and loss moduli remained at 1221.8 ± 342.6 Pa and 113.1 ± 18.5 Pa for DSPG/sLip-P407[15%], 1023.6 ± 332.1 Pa and 98.2 ± 34.7 Pa for DOTAP/sLip-P407[15%], respectively at body temperature, with a gelation temperature of 27-28°C.

**Fig. 3.**
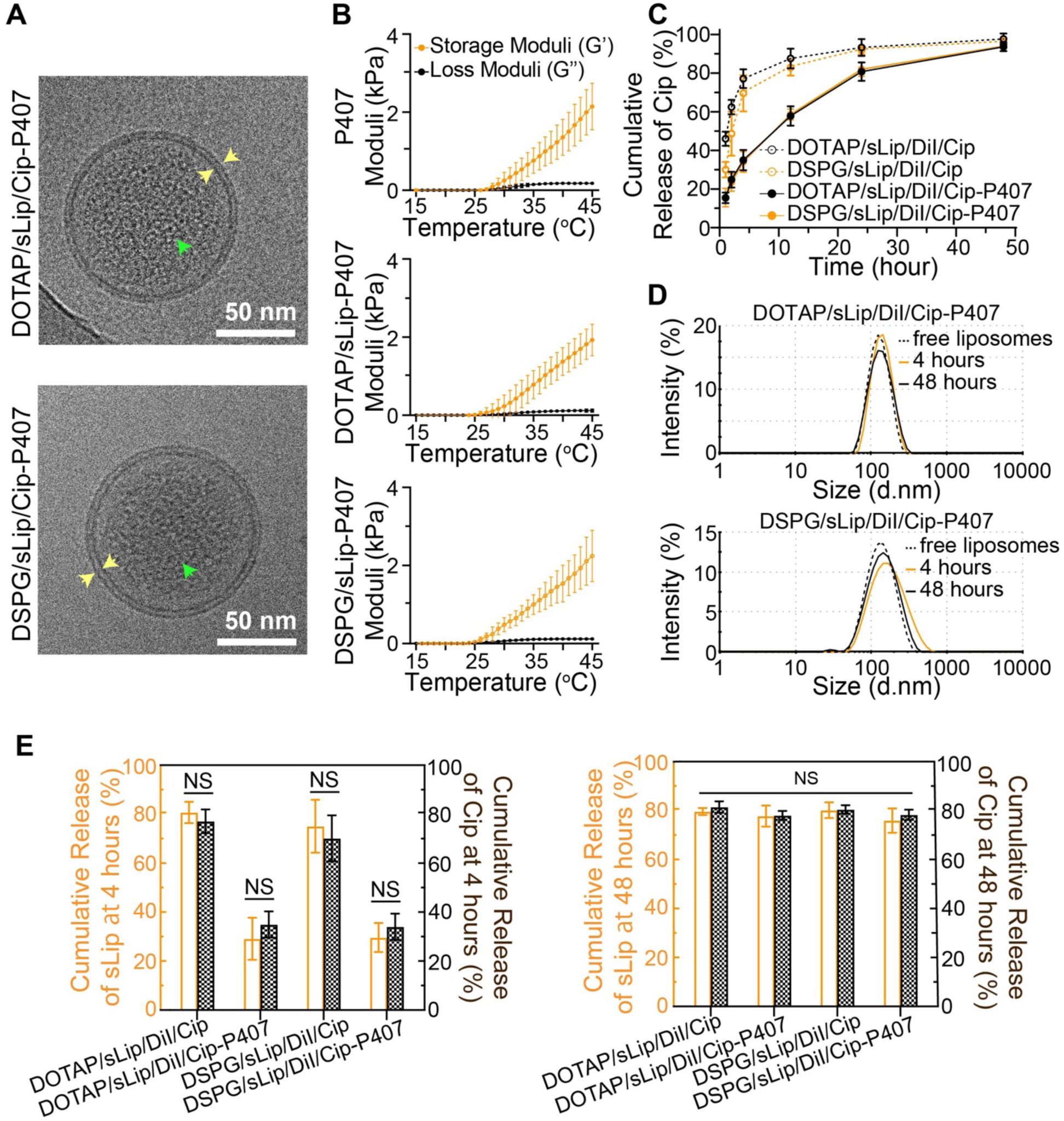
Stability, gelation, and in vitro release profiles of the liposome-containing hydrogel formulations. (**A**) Cryogenic transmission electron microscope (Cryo-EM) images of DOTAP/sLip/Cip-P407 and DSPG/sLip/Cip-P407. The yellow arrows indicate the lipid bilayer membranes, and the green arrows indicate the Cip crystals. Scale bars, 50 nm. (**B**) Rheology of the hydrogel formulations, including P407, DOTAP/sLip-P407, and DSPG/sLip-P407 (n = 3). (**C**) Cumulative release of Cip under infinite sink conditions over 48 hours (n = 3). (**D**) Size distributions of the liposomes released from the formulations tested in (C), at 4 and 48 hours; free liposomes (dotted lines) are free of P407 (n =3). (**E**) Release of intact liposomes (sLip, orange) and Cip (black) from the formulations tested in (C), at 4 and 48 hours (n = 3). All formulations contain 0.5 mg Cip; in all formulations containing P407, its concentration is 15% (w/v). Data are mean ± s.d. See Fig. S8 for the rheology of formulations containing 10% (w/v) or 20% (w/v) P407.

Using cryogenic electron microscopy (Cryo-EM), we confirmed that DSPG/sLip and DOTAP/sLip remain intact in the hydrogel after 24 hours of storage at room temperature (Fig. 3A). Their lipid bilayer membranes and the encapsulated Cip (in its crystal form due to dehydration) are clearly identifiable under Cryo-EM. The diameter of the liposomes was measured to be 112.1 ± 13.2 nm and 116.5 ± 15.1 nm for DSPG/sLip and DOTAP/sLip, respectively. That is smaller than the diameter obtained using DLS, likely due to the shrinkage caused by the liquid nitrogen freezing (as part of the Cryo-EM sample preparation). The lipid membranes exhibited a thickness of 5.0 ± 0.7 nm and 5.1 ± 0.8 nm for DSPG/sLip and DOTAP/sLip, respectively, and no rupture of the lipid membranes was observed.

The in vitro release kinetics was examined by placing the liposomes-containing hydrogel in a Transwell® insert and collecting the solution in the receiving chamber at 4 and 48 hours. The structural integrity of the liposomes upon release was analyzed with DLS (Fig. 3D). At 4 hours, particles in the receiving solution (i.e., 0.9% NaCl solution) demonstrate an average diameter of 134.1 ± 9.5 nm for DSPG/sLip/DiI/Cip-P407 and 136.0 ± 5.0 nm for DOTAP/sLip/DiI/Cip-P407, which became 131.2 ± 5.1 nm and 140.2 ± 27.4 nm, respectively, at 48 hours. The average particle sizes are close to the free liposomes (dotted line in Fig. 3D), i.e., 130.7 ± 4.7 nm for DSPG/sLip/DiI/Cip and 132.5 ± 4.3 nm for DOTAP/sLip/DiI/Cip (Table S1), indicating that the liposomes remain intact during the release from the P407 hydrogel.

The in vitro release kinetics of liposomes were obtained by measuring the fluorescence intensity (from DiI) of the receiving chamber solution (orange columns in Fig. 3E). Note that disintegrated liposomes do not emit fluorescence because free DiI is insoluble in saline, and the red fluorescence thus reflects intact liposomes. At 4 hours, 29.6 ± 5.9% and 29.1 ± 8.6% of the liposomes in DSPG/sLip/DiI/Cip-P407 and DOTAP/sLip/DiI/Cip-P407 were released, respectively, which increased to 91.1 ± 6.0% and 93.3 ± 5.1% at 48 hours. The surface charge or chemistry of the liposomes did not affect their release kinetics from the P407 hydrogel. The in vitro release kinetics of Cip were analyzed using High-Performance Liquid Chromatography (HPLC) (Fig. 3C). The results closely match those of the liposomes (Fig. 3E), indicating undetectable leakage of the encapsulated drug from the liposomes during their diffusion through the hydrogel matrix. The hydrogel slows the overall release kinetics of Cip for the first 4 hours, from 70.2 ± 9.3% to 34.0 ± 5.4% for DSPG/sLip/DiI/Cip and from 77.1 ± 4.9% to 34.9 ± 5.3% for DOTAP/sLip/DiI/Cip, but not by 48 hours, when 94.0 ± 2.5% and 93.6 ± 2.2% Cip is released from DSPG/sLip/DiI/Cip-P407 and DOTAP/sLip/DiI/Cip-P407, respectively.

While in vitro release studies are standard for teasing out the transport resistance within the delivery vehicle, i.e., hydrogel, the results may not represent the overall rate of drug delivery. In the context of transtympanic delivery, the overall kinetics is dominated by the most significant transport resistance, i.e., the permeation resistance across intact TM (*53, 63*). As such, results from ex vivo permeation experiments, as described below, better reflect the overall rate of drug diffusion (*53, 64*).

### The liposome-containing hydrogel enables transtympanic antibiotic delivery ex vivo

Using intact TM excised from animals with AOM, we assessed the ex vivo permeability of Cip achieved by the hydrogel formulations. Despite the additional transport resistance exerted by the hydrogel matrix, DSPG/sLip/Cip-P407 demonstrated the second highest transtympanic delivery rate in infected ears among all formulations tested (Fig. 2C). Cumulatively over 48 hours, 6.0 ± 2.2% of the Cip permeated across the TM from DSPG/sLip/Cip-P407, which is 3.8-fold that achieved by DOTAP/sLip/Cip-P407 (*P* = 0.0170) and 5-fold Cip-P407 (*P* = 0.0097). This result also corroborates the intactness of the excised TM, proving that therapeutics did not simply leak through the TM via perforations. Had perforations occurred, Cip should diffuse faster than liposomes, leading to the greatest flux for unencapsulated Cip, i.e., the Cip-P407 formulation. That was not the case. In fact, Cip-P407 (Fig. 2C, blue) achieved the lowest flux.

Notably, mixing (instead of encapsulating) Cip with DSPG/sLip led to no permeation enhancement effect, with 1.6 ± 0.4% (*P* = 0.0177 v.s. DSPG/sLip/Cip-P407) Cip permeating across infected TM in 48 hours (purple line in Fig. 2C). This result confirms that the DSPG liposomes do not function as simple chemical permeation enhancers (CPE), and that DSPG/sLip cross the TM much faster than Cip, pointing to active transport instead of passive diffusion.

While the ex vivo permeability measurement confirmed the viability of leveraging innate immune response to enhance the transtympanic permeation of therapeutics, it was performed on isolated auditory bullae. The immune system (besides the chemotaxis signals that may have leached into the receiving chamber), including the key immune cells and complement proteins, is missing or deactivating over the course of the experiment. Therefore, we employed an in vivo AOM disease model to assess the overall delivery efficiency, as detailed below.

### The liposome-containing hydrogel cures AOM in chinchillas in vivo

Prior to performing the in vivo efficacy testing, we assessed the cytotoxicity of the formulations and their constituents in human dermal fibroblasts (hFBs) (Fig. S9, A and C) and pheochromocytoma cells (PC12, frequently used to test neurotoxicity) (Fig. S9, B and D). While P407 hydrogel alone shows minimal cytotoxicity, the addition of 0.5 mg Cip leads to considerable cytotoxicity on day 3, with cell viability of 45.9 ± 5.3% for hFBs and 12.1 ± 0.2% for PC-12. DSPG/sLip/Cip-P407 reduced the cytotoxicity elicited by free Cip, increasing the 3-day cell viability to 88.3 ± 7.4% and 20.3 ± 6.8% for hFBs and PC12 cells, respectively.

The in vivo efficacy testing starts with the establishment of AOM in chinchillas by direct inoculation, as described above. To confirm the infection, evaluate treatment efficacy, and quantify the in vivo pharmacokinetics, MEF was aspirated through the auditory bullae following a schedule shown in Fig. 4A. The criterion for a successful establishment of infection is that 3 days post-inoculation, the concentration in the MEF should be > 10^6^ colony-forming unit (CFU)/ml. In infected animals that did not receive any treatment, the bacterial concentration of > 10^5^ CFU/ml was maintained throughout the 7-day course of monitoring (Fig. 4C). In treated animals, 200 µL test formulation which contained 0.5 mg ciprofloxacin, was deposited onto each TM through the external auditory canal. The TMs were closely monitored using otomicroscope throughout the in vivo experiment, following a procedure reported by Kohane, et al (*52*). Animals with perforated TM, identified via otomicroscope or during the dissection at the end of each experiment, were excluded from the reported results.

**Fig. 4.**
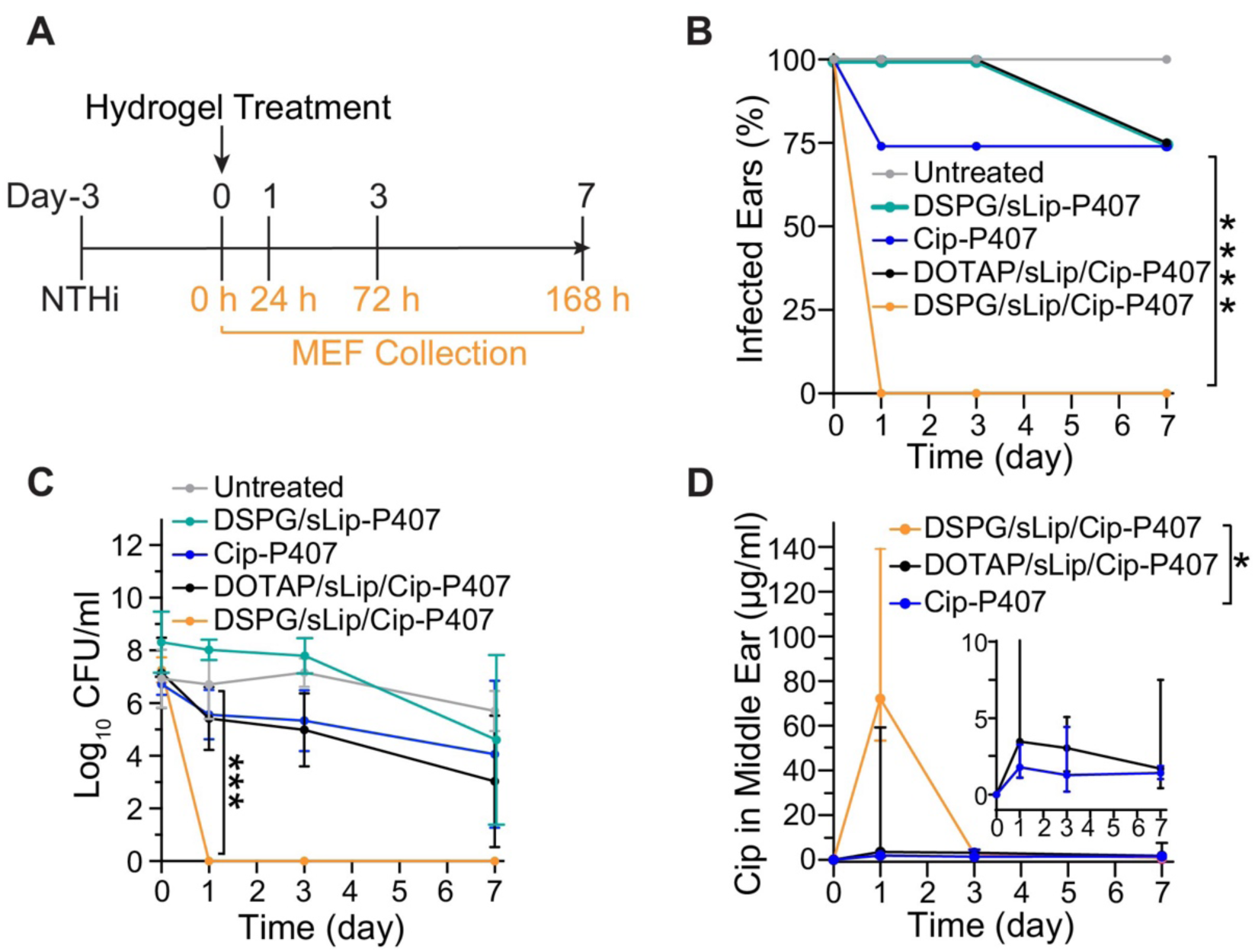
In vivo efficacy and pharmacokinetics of DSPG/sLip/Cip-P407. (**A**) Diagram of the in vivo experimental procedure. (**B**) Percentage of animals with AOM from NTHi before and after receiving DSPG/sLip-P407, Cip-P407, DOTAP/sLip/Cip-P407, and DSPG/sLip/Cip-P407. Day 0 reflects the status immediately before the administration of therapeutics (n = 4). \*\*\*\**P* < 0.0001 by Fisher’s exact test. (**C**) Time courses of bacterial CFU from MEF aspirated from chinchillas with AOM treated with DSPG/sLip-P407, Cip-P407, DOTAP/sLip/Cip-P407, and DSPG/sLip/Cip-P407 (n = 4). (Log_10_ CFU is set to 0 instead of minus infinity for the purpose of this illustration.) \*\*\**P* = 0.0001 by one-way analysis of variance (ANOVA) with Turkey’s test. (**D**) Concentration of Cip over time in the MEF of the same animals as in (B) and (C) (n = 4). Inset is the magnified drug concentration range of 0 to 10 µg/ml. \**P* = 0.0142 by nonparametric Kruskal-Wallis multiple comparisons test. Each formulation contains 0.5 mg Cip, 2 mg lipids, and 15% (w/v) P407, unless otherwise specified. Data are medians with the 25th and 75th percentiles in parentheses. See Fig. S7A for the corresponding cumulative transtympanic permeation (%) of Cip over time, and Fig. S7D for a comparison of the ex vivo and in vivo across TM ratios of the cumulative Cip at 24 hours. See Table S2 for the normality test of data in (C) and (D) with the Shapiro-Wilk Test.

In infected animals that received DOTAP/sLip/Cip-P407 or Cip-P407, NTHi was eradicated in 25% of the treated ears by day 7 (n = 4, Fig. 4B). The titre of pathogens in the MEF remained high (Fig. 4C), at 10^3.0^ ± 10^2.5^ CFU/ml for DOTAP/sLip/Cip-P407 (*P* = 0.2482 v.s. untreated), and 10^4.1^ ± 10^2.8^ CFU/ml for Cip-P407 (*P* = 0.6270 v.s. untreated). An antibiotic-free formulation, i.e., DSPG/sLip-P407, was also tested to assess if the innate immune response alone can clear AOM. This formulation achieved a 25% cure rate, with a MEF pathogen titre of 10^4.6^ ± 10^3.3^ CFU/ml (*P* = 0.9563 v.s. untreated). In contrast, NTHi was eradicated in 100% of the animals received DSPG/sLip/Cip-P407. That complete cure was achieved within the first 24 hours of the treatment, as indicated by the zero CFU in the MEF (Fig. 4C) (*P* = 0.0001 v.s. untreated). The sterile auditory bullae were maintained until the end of the experiment on day 7.

The 100% cure rate was a direct result of the large fraction of ciprofloxacin delivered into the middle ear by DSPG/sLip/Cip-P407 (Fig. 4D). As quantified using HPLC, a peak Cip concentration of 72.0 µg/ml (median; IQR = 53.3-139.1 µg/ml) was achieved by DSPG/sLip/Cip-P407, representing up to 600-fold the reported minimum inhibitory concentration (MIC) of Cip against NTHi (*65*). Assuming a middle ear volume of 1 ml for chinchillas (*66*), 14.4% (median; IQR = 10.7%-27.8%) of the total Cip contained in DSPG/sLip/Cip-P407 was delivered across TM within 24 hours (Fig. S7A). That delivery efficiency (at Day 1) was 21-fold that achieved by DOTAP/sLip/Cip-P407, i.e., 0.7% (peak median concentration = 3.5 µg/ml; IQR = 1.1-59.2 µg/ml; *P* = 0.0955 v.s. DSPG/sLip/Cip-P407), and 36-fold that achieved by Cip-P407, i.e., 0.4% (peak median concentration = 1.8 µg/ml; IQR = 1.1-3.3 µg/ml; *P* = 0.0142 v.s. DSPG/sLip/Cip-P407) (Fig. S7A).

The delivery efficiency of DSPG/sLip/Cip-P407 can be further increased upon exacerbation of the native immune response. We used a coinfection animal model (where animals were simultaneously infected with NTHi and *Streptococcus Pneumoniae*) to achieve that heightened immune response (Supplementary Methods). The treatment with 200 µL DSPG/sLip/Cip-P407 led to a median Cip concentration of 262.4 ± 211.7 µg/ml in the MEF after 24 hours (Fig. S7B), representing a median delivery efficiency of 52.5 ± 42.3%. The experiment was terminated on Day 1 due to the development of pneumonia in the infected animals. As no perforation was observed on the TM of the treated animals, we believe the higher delivery efficiency was a result of the heightened immune reactions. A direct connection between the transtympanic delivery efficiency and the complement cascade is detailed in the section below.

### The complement cascade is key to the in vivo transtympanic drug delivery and successful AOM treatment

We designed the hydroxyl-bearing liposomes to interact with the complement cascade, which enhances their internalization by neutrophils and transportation to the site of infection, i.e., the middle ear (Scheme. 1). To illustrate the key role played by the complement cascade in vivo, we depleted the complement protein fragments from the systemic circulation of the infected animals and demonstrated that the transtympanic delivery was no longer effective.

The complement proteins were depleted by treating infected animals with Cobra Venom Factor (CVF), a structural analogue of C3, and a non-toxic protein purified from cobra venom (*31*). The treatment with CVF has been reported to deplete complement proteins and achieve an anti-complement effect by its continuous and sustained activation of C3 (*31, 32*). CVF was injected at a dose of 500 mg/kg body weight (via femoral vein) 24 hours prior to the hydrogel treatment (Fig. 5A). The C3b and its iC3b fragments in the MEF and serum of the treated animals were undetectable via western blot on Day 1 (Fig. 5B) and remained nearly undetectable for up to 3 days post-hydrogel-treatment.

**Fig. 5.**
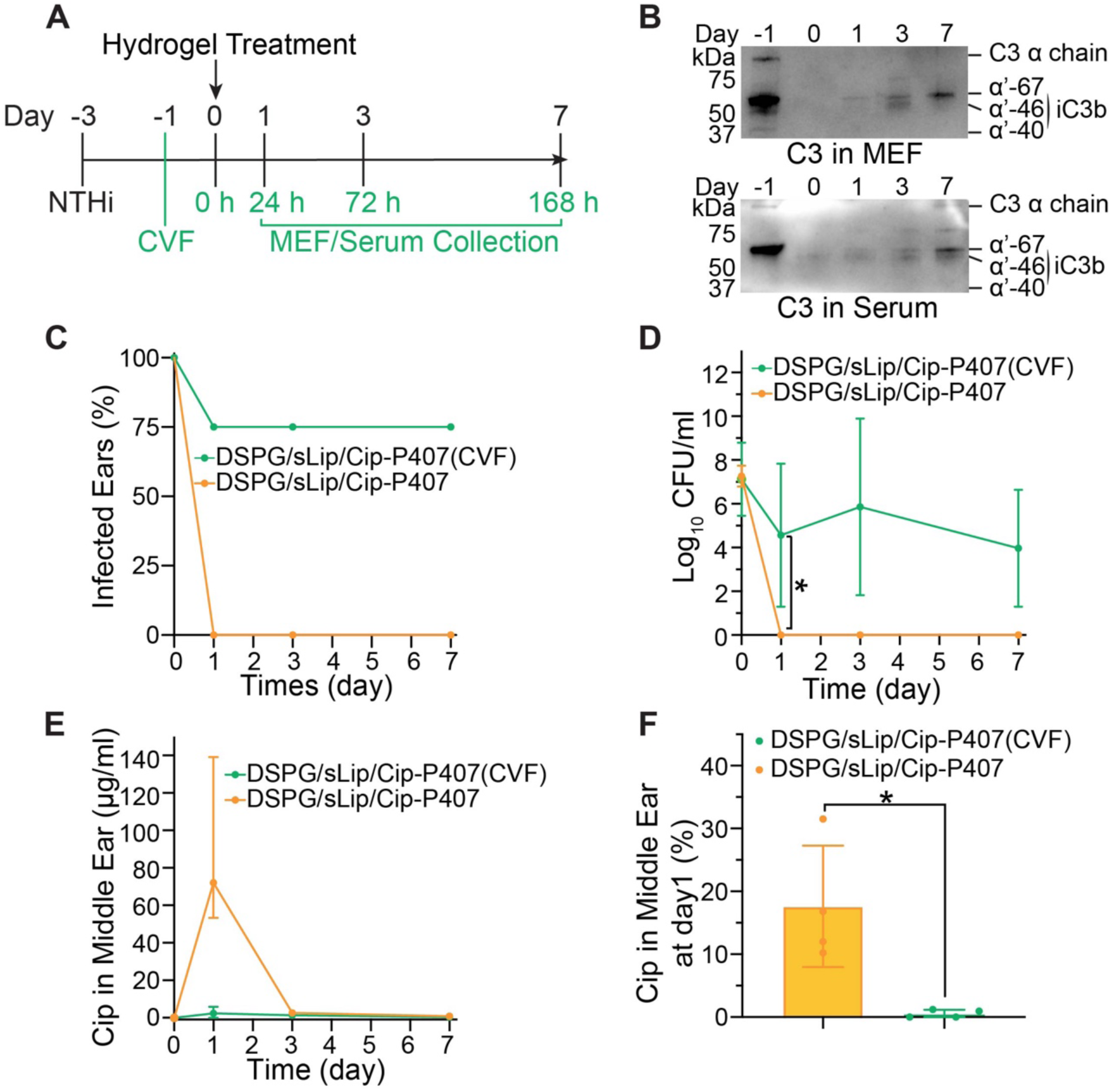
In vivo efficacy of DSPG/sLip/Cip-P407 in C3b-depleted animals. (**A**) Diagram of the in vivo experimental procedure to deplete C3b fragments, using Cobra Venom Factor (CVF), in infected chinchillas. (**B**) Western blot showing the levels of C3b fragments over time in the MEF and serum sampled from the CVF-treated chinchillas (same in (C-F)). (**C**) Percentage of animals with AOM after receiving DSPG/sLip/Cip-P407, with and without the CVF treatment (n = 4). (**D**) Time course of the bacterial CFU from the MEF of the same animals as in (B-C) (n = 4). \**P* = 0.0313 by two-tailed unpaired Student’s t-test. (**E**) Concentration of Cip over time in the MEF of the same animals as in (B-D) (n = 4). (**F**) Cumulative permeation of Cip across the TM within the first 24 hours of the hydrogel treatment (in terms of a percentage of the total Cip applied) (n = 4). \*\**P* = 0.0124 by two-tailed unpaired Student’s t-test. The data on the animals received DSPG/sLip/Cip-P407 and no CVF were replotted from Fig. 4 as a point of comparison. Data are medians with the 25th and 75th percentiles in parentheses. See Fig. S7A for the corresponding cumulative transtympanic permeation (%) of Cip over time.

In the animals that received CVF, DSPG/sLip/Cip-P407 eradicated NTHi in 25% of the infected ears by day 7 (Fig. 5C), with an NTHi titre of 10^4.0^ ± 10^2.7^ CFU/ml in the MEF on day 7 (Fig. 5D). The results for animals treated with DSPG/sLip/Cip-P407 and no CVF (Fig. 4) were replotted in Fig. 5 to serve as a point of comparison. The Cip concentration in the MEF of the CVF-treated animals was 2.7 ± 3.2 µg/ml, captured on Day 1 (Fig. 5E and Fig. S7A), corresponding to 0.5 ± 0.6% of the total Cip in the formulation. As such, there is a 33-fold difference in MEF drug concentration when the neutrophil chemotaxis and complement cascade are uninhibited versus inhibited (*P* = 0.0124; Fig. 5F).

Interestingly, CVF is also known to deplete C5a (*67*), a complement component and a neutrophil chemoattractant released from tissues near the site of infection (*68*). C5a plays a critical role in the local recruitment of neutrophils as it is known to override the signals by chemoattractants released by intermediary sites, such as the endothelium (*69*), which has been considered a dominating mechanism to trigger neutrophil chemotaxis (*70*). Therefore, the treatment with CVF likely also reduces the local recruitment of neutrophils, which further contributes to the reduced efficacy. Taken together, results from the CVF treatment confirm that the complement cascade is essential for achieving therapeutic antibiotic levels in the middle ear.

### The liposome-containing hydrogel is biocompatible in the ear

TM harvested from infected but untreated animals exhibit an average thickness of 57.4 ± 11.2 µm, 5.8-fold that of the pristine TM (9.9 ± 2.0 µm) (Fig. 6A). Acute inflammation was observed with diffuse edema and dense infiltration by inflammatory cells. TM excised from the animals infected and treated with DSPG/sLip/Cip-P407 for 7 days appear nearly identical to the pristine ones, with an average thickness of 12.3 ± 1.4 µm and no tissue injuries, necrosis, or inflammatory cells (Fig. 6A). By comparison, animals that received Cip-P407 (40.3 ± 12.0 µm), DOTAP/sLip/Cip-P407 (46.0 ± 7.4 µm), DSPG/sLip-P407 (58.5 ± 13.0 µm), or CVF injection prior to the DSPG/sLip/Cip-P407 treatment (50.1 ± 12.1 µm) demonstrate considerable edema and inflammation in their TMs (Fig. 6A). The mucosal layer that lines the middle ear bullae remained unchanged after the treatment with DSPG/sLip/Cip-P407 for 7 days compared to the pristine bullae (Fig. S10), confirming the biocompatibility of the treatment in the ear.

**Fig. 6.**
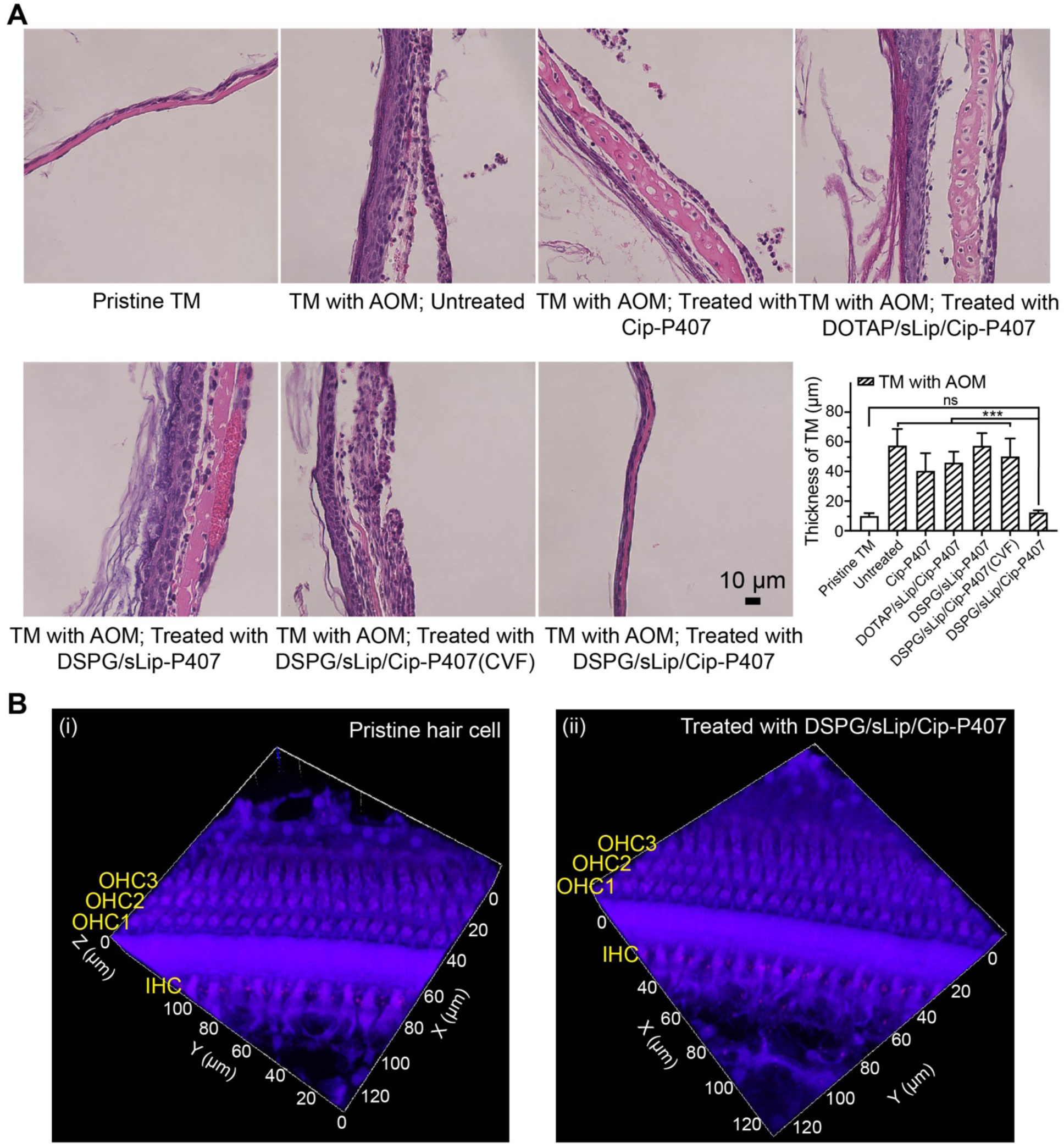
Biocompatibility of the DSPG/sLip/Cip-P407 formulation in the ear. (**A**) Representative photomicrographs of hematoxylin and eosin (H&E)-stained cross-sections of healthy and untreated TM, TM with AOM and left untreated, and TM with AOM and treated for 7 days using Cip-P407, DOTAP/sLip/Cip-P407, DSPG/sLip-P407, CVF (on Day -1) followed by treatment using DSPG/sLip/Cip-P407, or DSPG/sLip/Cip-P407. Labeled on each panel is the TM’s average thickness ± s.d. (n = 4). Scale bar, 10 µm. \*\*\**P* < 0.001 by one-way analysis of variance (ANOVA) with Turkey’s test. (**B**) Chinchilla inner ear hair cells, pristine (i) or those collected from healthy animals treated with DSPG/sLip/Cip-P407 for 7 days (ii), stained with Phalloidin (red) and Hoechst (blue). Outer hair cell, OHC; inner hair cell, IHC. The concentration of P407 is 15% (w/v); the amount of ciprofloxacin in formulation is 0.5 mg.

Additionally, the biocompatibility of DSPG/sLip/Cip-P407 in the ear was evaluated based on its effects on inner ear hair cells and hearing sensitivity. As shown in Fig. 6B, the morphology and arrangement of the three layers of outer hair cells (OHC) and one layer of inner hair cells (IHC) look identical to the pristine ones (Fig. 6B-i) after 7 days of treatment with the DSPG/sLip/Cip-P407 formulation (Fig. 6B-ii). The effect of DSPG/sLip/Cip-P407 on hearing sensitivity was assessed via the auditory brainstem response (ABR) of healthy chinchillas before and after receiving the formulation. Placement of 200 µl DSPG/sLip/Cip-P407 on the TM did not cause a detectable shift of the ABR threshold (Fig. S11), indicating unchanged hearing sensitivity as detailed in previous studies (*18*).

## DISCUSSION

We have developed a fresh concept in transtympanic drug delivery, which led to a formulation that cures acute otitis media (AOM) by targeting antibiotics directly to the middle ear. The formulation eradicates AOM in 100% of the treated animals within 24 hours, using 0.5 mg antibiotic. Furthermore, it was designed to allow a pediatrician to deliver a single dose of antibiotics to TM that provides an entire course of treatment, reducing pediatric antibiotic usage and improving patient compliance.

The TM is a tri-layer membrane, where the outmost layer is a keratinized stratified squamous epithelium continuous with the skin of the external auditory canal (*71*). It is one of the most impermeable biological barriers, distinguishing it from other tissues in the ear, including the middle ear mucosa, round window membrane, and tissues that do not contain stratified squamous epithelial cells (*72*). Like in skin, the barrier function of the TM predominantly resides within the stratum corneum layer (*43*). The transtympanic delivery of therapeutics has been a longstanding challenge and the focus of many recent studies on AOM treatment (*18, 19, 73–76*). While the current state-of-the-art approaches often rely on the structural disruption of TM, e.g., via tympanostomy or molecular disruption using chemical permeation enhancers, the reported strategy is conceptually distinct as it leverages the innate immune response of children with AOM to overcome the impermeable TM. Leveraging the innate neutrophils as drug carriers also leads to a targeted therapeutic effect, as neutrophils migrate to the site of infection in response to invading pathogens. Although neutrophils are known to infiltrate the middle ear mucosa (*42, 77*), their ability to penetrate the TM is surprising because the TM is known to be impermeable to even small molecules, like Helium gas (*78*) (with a molecular diameter of 2.6 Å). The paradigm of hitchhiking on immune cells to overcome the stratum corneum could be broadly applicable to treating diseases beyond AOM, such as topical and on-demand treatments for skin disorders. External biological barriers, such as bacterial biofilms that often cause chronic or untreatable infections, are also potential targets for the penetration effects via immune cell hitchhiking (*79, 80*).

The design also overcomes a challenge in harnessing neutrophils as drug delivery vehicles, i.e., the short lifetime of neutrophils (*t*½ < 24 hours) (*81, 82*). As most existing systems rely on harvesting neutrophils ex vivo for drug loading, their short lifetime, combined with the laborious procedures of neutrophil isolation, purification, drug loading, and re-infusion into the bloodstream, limits the success of such treatments (*14, 83*). While considerable efforts have been made to design nanoparticles that hitchhike on circulating neutrophils, the systemically injected particles have been shown to be cleared quickly by the reticuloendothelial system, leading to low delivery efficiency (*84*). We overcome these challenges by delivering the neutrophil-hitchhiking liposomes locally and topically, escaping systemic clearance while avoiding off-target effects. Neutrophil levels throughout an episode of AOM are a critical consideration during the bench-to-bedside translation of this technology. Recent clinical studies indicate that the neutrophil levels in the ear, nose, and throat (ENT) systems are consistently 2-4 orders of magnitude greater in children with AOM than in healthy children (*85*). While these studies lend strong support to the potential of the reported drug delivery system in addressing pediatric AOM, we emphasize that significant further investigations (e.g., on the intervention timing) are required before expanding this strategy for clinical treatment.

To load antibiotics into innate neutrophils, we designed hydroxylated liposomes that are preferentially opsonized by the complement protein (C3b) fragments for enhanced neutrophil internalization. The preferential opsonization was illustrated using western blot. The effective hitchhiking of hydroxylated liposomes on neutrophils was demonstrated in a mouse model using flow cytometry. The traversing of the liposome-bearing neutrophils across intact chinchilla TM was visualized using immunocytochemistry analysis. The key role of the complement cascade was further illustrated in vivo by depleting the complement protein fragments in infected animals and observing that the hydroxylated liposomes no longer enhance transtympanic permeation or eradicate AOM. The mammalian stratum corneum comprises broad lamellar sheets of negatively charged lipids, including ceramides (40%), free fatty acids (25%), cholesterol (25%) and cholesteryl sulfate (10%) (*86*). Therefore, positively charged liposomes, such as DOTAP/sLip, have been considered favorable for enhancing transtympanic permeation via electrostatic interactions (*87, 88*), which is consistent with our ex vivo results on the permeability of healthy TM. However, during AOM, the hydroxylated liposome DSPG/sLip exhibits a superior transtympanic permeability than the positively charged liposomes, reversing the trend observed in healthy ears. Notably, the in vivo transtympanic delivery efficiency achieved by DSPG/sLip was 7-fold its ex vivo efficiency after 24 h treatment, which is the opposite of what has been observed in previous transtympanic delivery studies (*18, 19, 73, 74*). That greater delivery efficiency in vivo hints at the enabling role played by the innate immune system, which is only present in live animals. Furthermore, we unveiled a positive correlation between the transtympanic delivery efficiency and the severity of the infection, pointing to a smart and self-regulated treatment.

The formulation comprises a temperature-responsive hydrogel that is easy to apply under room temperature and gels quickly upon contacting the warm TM, streamlining the treatment of young patients who are most prone to AOM. A single dose can be applied by a pediatrician at the time of diagnosis, which delivers the entire course of the regimen to cure the infection. Poloxamer 407 (P407) was used to formulate the temperature-responsive hydrogel. Using Cryo-EM, rheology, and dynamic light scattering (DLS), we confirmed that the liposomes remain intact upon loading into or release from the P407 hydrogel, and the liposomes do not disrupt the reverse thermal gelation of P407. Furthermore, P407 has been approved by the FDA as a polymeric drug carrier, which, combined with the many FDA-approved liposomal formulations, promises rapid progression of the reported treatment toward clinical adoption. While ciprofloxacin was chosen based on its broad spectrum of activity against relevant bacteria and its effectiveness against AOM in children with myringotomy tubes, liposomes are known for their encapsulation versatility. The delivery system is likely compatible with other drugs. In addition, liposomes are often used to deliver drugs with poor bioavailability and/or solubility, which could be leveraged to broaden the range of antimicrobials applicable in AOM treatment.

Our future research will focus on understanding the correlations between particle size and the efficiencies of neutrophil uptake, transtympanic delivery, and infection eradication. While liposomes with a diameter of ∼100 nm were used in this study, based on the reported preference of neutrophils for nanometer-scale particles to micrometer ones (*89, 90*), there is room for further optimization. In addition, while the current study focuses on the effect of neutrophils in transtympanic delivery, it is possible that other native immune cells and responses contributed to the overall efficacy, which will be dissected in detail in the future. While *Chinchilla lanigera* is one of the most prevalent animal models for AOM studies, the safety and efficacy results of which have supported human clinical trial applications (*91*), we emphasize that the TM thickness and immune responses can differ significantly between the two species. These factors could limit the future success of the reported delivery system during clinical translation. The strategy presented here leverages the active transport of neutrophils instead of passive diffusion (e.g., for chemical penetration enhancers). While the thicker human TM is anticipated to lead to slower passive diffusion, that effect is unlikely applicable to active transport. In fact, existing reports on immune cells crossing biological barriers (like the blood-brain barrier) revealed little evidence for such effects (*92*). The limited number of comparative studies of the human immune responses versus those of chinchillas further reduced the confidence in any prediction made regarding the ultimate efficacy in human patients.

## MATERIALS AND METHODS

### Study design

We aimed to develop a noninvasive, transtympanic drug delivery platform for OM. We focused on developing and evaluating formulations containing liposomes and an antibiotic. The experiments compared the effect of the polymer matrix and incorporation of liposomes on TM permeability and OM cure rate. During ex vivo experiments, data collection was stopped after 48 hours due to microbial growth on harvested TM, whereas, during in vivo experiments, data collection was stopped after 7 days because OM would either be cleared or cause the animal severe illness that required euthanasia. In vivo experiments were blinded. The information regarding whether the animal was infected or healthy and regarding which formulation was administered was kept by two separate research groups and not shared until all data analyses were done. Animals were assigned treatment groups at random.

### Animal maintenance and cells

Healthy adult chinchillas weighing 300 to 400 g were procured from the Kuby Kritters (Akron, USA) (NY). Healthy male BALB/C mice aged 4-6 weeks were from the Jackson Laboratory (NY). All animals were raised and cared for according to the animal experimentation protocols approved by the Institutional Animal Care and Use Committee (IACUC) in conjunction with the Animal Use Guidelines established by the Cornell University Center of Animal Resources and Education (CARE).

Human dermal fibroblast cells (hFBs; PCS-201-012) and PC12 cells (CRL-1721) were purchased from the American Type Culture Collection company (ATCC) and cultured in the recommended medium under 5% CO2 atmosphere according to ATCC instructions.

### Preparation and characterization of the protein corona

Middle ear fluid (MEF) and serum were sampled from NTHi-infected chinchillas (at 72 hours post inoculation). Liposomes (containing 1 mg lipid; 100 µl) were incubated with an equal volume of MEF or serum at 37°C for 1 hour to form protein corona and subsequently mixed with 1 ml chilled PBS. The mixture was centrifuged at 12000 g for 30 min at 4°C. The supernatant was discarded, and precipitate resuspended with 1 ml chilled PBS. The purification steps of the precipitate were repeated three more times, and in the final step, the precipitate was suspended in PBS and subsequently mixed with an equal volume of 2X Sample Buffer containing 0.55 M β-mercaptoethanol. To prepare for SDS-PAGE, samples were heated to 95 °C and incubated for 5 minutes. 10 µl of each as-prepared samples were loaded into a 4-20%-gradient gel, along with the pre-stained protein standards. The electrophoresis occurred under a constant current of 20 mA. Electroblotting (by placing the gel in direct contact with a piece of 0.45-µm nitrocellulose membrane) occurred under a constant current of 200 mV and over 70 minutes. The primary complement C3 polyclonal antibody and HRP-conjugated secondary antibody were used to detect the target C3 protein fragments. The levels of C3 protein fragments in the serum and MEF of infected animals were characterized using the same method.

### Hydrogel formation

P407 hydrogel formulations with different polymer concentrations (10, 15, 20% (w/v)) were prepared by dissolving P407 powder in deionized water or liposomes solution. Linear oscillatory shear rheology measurements were performed at 100 rad^-1^, 1% strain, and 1°C min^-1^. The changes in storage moduli (G’) and loss moduli (G’’) over temperatures from 15°C to 45°C were recorded with triplicates. The gelation temperature was determined as the temperature at which G’ became greater than G’’. Cryogenic transmission electron microscopy (Cryo-EM) was performed to analyze the stability of liposomes in the hydrogel. Applied the DOTAP and DSPG liposomes (3 µl, diluted ten-time with deionized water) to a support structure (grid), reduced its dimension to a layer that was as thin as possible (∼100-800 Å), and then froze this layer with liquid nitrogen to prevent the water from crystallizing before Cryo-EM imaging.

### Isolation of Mouse Neutrophils and ex vivo cell uptake evaluation

Obtaining a sufficient quantity of neutrophils from the peripheral blood of mice is challenging (*48, 49*); therefore, neutrophils were isolated from bone marrow. Male BALB/c mice were euthanized and immersed in 70% ethanol for 10 min to ensure full sterilization. The femur and tibia, along with attached muscle tissue, were carefully separated, followed by sterilization of the collected bones using 70% ethanol and subsequent washing with pre-chilled buffer (0.5% BSA and 2 mM EDTA in PBS, pH 7.2). Bone marrow cells were harvested by thorough flushing of the bone cavity with the aforementioned buffer and subsequent centrifugation at 400 g, 4°C for 5 minutes. The resulting pellet was suspended in 10 ml of 0.2% sterile NaCl solution to lyse the red blood cells, with osmotic pressure restored by mixing with 10 ml of 1.6% sterile NaCl solution after 30 seconds. The cell suspension was filtered through a 70 µm sterile cell strainer and centrifuged at 400 g, 4°C for 5 min. The pellet was suspended with 1 ml PBS for subsequent purification. In a 15 ml sterile tube, 5 ml Histopaque®-1077 (1.077 g/ml), 1 ml bone marrow cell suspension, and 5 ml Histopaque®-1119 (1.119 g/ml) were carefully layered from the bottom to the top without agitation. Neutrophils were collected by centrifugation at 1000 g, 4°C for 30 min, and washed twice with RPMI-1640 medium. To assess the purity of the isolated neutrophils, cells were stained with Anti-Mouse CD11b Antibody (APC (Allophycocyanin)) and Anti-Ly-6G Antibody (BV421 (Brilliant Violet reg 421)) at 0.2 µg of 1:1 mixed antibodies per 10^6^ cells in 100 µl PBS on ice. The purity of neutrophils was approximately 70% as confirmed by flow cytometry.

The isolated neutrophils were aliquoted to 10^6^ in 100 µl PBS (supplemented with 10% mouse serum). DiO-labeled liposomes were incubated with BALB/c mouse serum with 1:1 at 37°C for 1 hour to form protein corona. The opsonized liposomes (10 µM lipid) were mixed with 10^6^ neutrophils in 0.1 ml PBS and incubated at 37°C for 1 hour or 4 hours. Cell pellets were collected using a centrifuge (5425R, Eppendorf) and washed with PBS twice to fully remove unreacted liposomes. A total amount of 0.2 µg of antibodies, i.e., Anti-Mouse CD11b Antibody (APC (Allophycocyanin)) and Anti-Ly-6G Antibody (BV421 (Brilliant Violet reg 421)), mixed with a 1:1 ratio, was incubated with 10^6^ neutrophils at 4°C for 1 hour. Cell pellets were washed with PBS and fixed with 4% paraformaldehyde (preserving the samples and avoiding any potential contamination during analysis) for subsequent analyses using Flow Cytometry (FACSymphony A3, BD Biosciences).

### Mouse in vivo neutrophil uptake studies

Adult healthy BALB/c mice were injected intravenously with DiO-labeled liposomes through the tail vein at a dose of 50 mg lipids per kg bodyweight. Peripheral blood samples were collected through the retro-orbital vein into tubes with pre-chilled PBS supplemented with 10 mM EDTA. Blood cell pellets were collected by centrifugation (5425R, Eppendorf) and subsequently mixed with mouse red blood cell lysis buffer with moderate vortex. Cell pellets were collected by centrifugation and washed twice with PBS. A total amount of 0.2 µg of antibodies, i.e., Anti-Mouse CD11b Antibody (APC (Allophycocyanin)) and Anti-Ly-6G Antibody (BV421 (Brilliant Violet reg 421)), mixed with a 1:1 ratio, was incubated with 10^6^ neutrophils at 4°C for 1 hour. Cell pellets were washed with PBS and fixed with 4% PFA (preserving the samples and avoiding any potential contamination during analysis) for subsequent analyses using Flow Cytometry (FACSymphony A3, BD Biosciences).

### In vitro release studies

The release kinetics of ciprofloxacin from formulations were characterized using a Transwell® diffusion assay. Transwell® membrane inserts with 8-μm pore size and 1.1-cm^2^ area (Costar) and 24-well plates (notched for use with cell culture inserts, Corning) were used as the donor and receiving chamber, respectively. 200 µl pre-warmed formulation was added into each membrane insert and incubated at 37°C for 10 min to form a solid hydrogel. Subsequently, each membrane insert was immersed into a well containing 2 ml pre-warmed PBS and incubated at 37°C. Saline in the receiving wells was sampled at 0.5, 1, 2, 4, 6, 12, 24, and 48 hours. Samples were mixed with an equal amount of methanol to ensure complete drug dissolution and chromatographically analyzed with High-Performance Liquid Chromatography (HPLC) to determine ciprofloxacin concentration (λ = 275 nm). Experiments were performed in triplicates.

### Ex vivo TM permeation

The cross-TM permeation rate of ciprofloxacin was determined with auditory bullae harvested from healthy and NTHi-infected chinchillas (infected for 48 hours), where a current greater than 50 µA under the alternating voltage of 100 mV was used as an indication of micro/macro-perforations. All formulations were applied into the bullae kept at 37°C and deposited onto the TM. The volume applied was 200 µl, translating to 0.5 mg of ciprofloxacin. Permeation of ciprofloxacin across TM into the receiving chamber was quantified using HPLC. Detailed information regarding TM harvesting, TM electrical resistance measurement, and ex vivo permeation experiment configuration can be found in our previous reports (*5, 18*).

### NTHi OM chinchilla model

All procedures and manipulations were performed in accordance with the approved protocol by the IACUC and the guidelines by Cornell University CARE. Isolates of NTHi grown to the mid-log phase were diluted in Hanks balanced salt solution (HBSS), and about 100 to 200 CFU in 200 µl were introduced directly into the auditory bullae using the well-established transbullar inoculation technique for inducing experimental otitis media in chinchillas (*93, 94*). Daily otomicroscopy were performed to determine the presence of fluid in the auditory bullae and signs of infection, including bulging TM. Once abnormality was identified, the middle ear cavity was accessed 48 to 72 hours later. TMs of the animals to receive the formulation were observed with the speculum of an otoscope, after which the liquid hydrogel was injected through the speculum using a soft 18-gauge, 1.75-inch catheter. A direct culture of middle ear was obtained with a calcium alginate swab and immediately streaked onto a chocolate agar plate. Middle ear fluid was obtained with an 18-gauge angiocatheter connected to an empty syringe, 10 to 20 µl of MEF were diluted 1:10 in HBSS, and eight serial 10-fold dilutions were prepared. Ten microliters of each dilution were plated onto the chocolate agar with duplicates. The lower limit of detection of viable organisms in MEF using this dilution series was 1 CFU/µl. Direct and indirect ear examination was performed every 1 to 2 days until the middle ear cultures were sterile. Note that the ex vivo and in vivo experimental protocols let the chinchillas develop AOM for different numbers of days post-inoculation, following established methods(*5, 53*). During ex vivo experiments, animals were euthanized and dissected once the symptoms of AOM (such as inflamed TM and recruitment of neutrophils to the ear) were established. That typically occurs two days after inoculation. During in vivo experiments, it is critical that animals establish an episode of AOM that cannot be cleared by their innate immune system, calling for a middle ear colony-forming unit (CFU) concentration beyond 10^6^ CFU/ml, which typically takes 3 days.

To study the interaction between liposomes and neutrophils, 200 µl DPSG/sLip/DiI-P407 or DOTAP/sLip/DiI-P407 were applied to the TM of the infected chinchillas through the external auditory canal. After 24 hours, the TMs and MEF were sampled and stained with Hoechst for the cell nucleus and anti-elastase antibody for neutrophil. Zeiss 710 confocal microscope was used for imaging.

### Hair cell and ABR measurements

Healthy chinchillas were treated with 200 µl DSPG/sLip/Cip-P407 formulation, administered onto their TMs through external ear canals, and sacrificed after 7 days of treatment. The inner ear cochlear hair cells were carefully dissected under a microscope and stained with Hoechst for the cell nucleus and Phalloidin (conjugated with CF568) for F-actin, respectively. Zeiss 710 confocal microscope and Keyence BZ-180 microscope were used for fluorescence and brightfield imaging, respectively.

ABR experiments were conducted using a custom-designed stimulus generation and measurement system built around the National Instruments software (LabVIEW) and hardware, as described in Supplementary Materials and Methods.

### Cytotoxicity evaluation

Two hundred microliters of each test formulation were deposited into an 8-μm Transwell® membrane insert (Costar), which was subsequently placed in a 24-well plate seeded with hFB or PC12 (American Type Culture Collection). After 1 or 3 days of incubation, cell viability was evaluated using the 3-(4,5-dimethyl2-yl)-5-(3-carboxymethoxyphenyl)-2-(4-sulfophenyl)-2H-tetrazolium inner salt (MTS) assay (for mitochondrial metabolic activities), i.e., CellTiter 96 AQueous One Solution Cell Proliferation Assay (Promega Corp.). The absorbance at 490 nm was recorded using a plate reader (Tecan launches Infinite® M1000 PRO) and used to quantify cell viability. Cell viability was confirmed using a LIVE/DEAD Viability/Cytotoxicity Kit (Molecular Probes, Invitrogen).

### Statistical analyses

For the in vivo experiments, a sample size of 4 is used, which is supported by prior publications in this field (*5, 19, 74*). Statistical analyses are conducted using GraphPad software (version 9.4.1, San Diego, CA). All statistical analyses used are indicated in the corresponding figure captions. The Shapiro–Wilk test is applied to test the normality of the data (*95*), and the Shapiro– Wilk test results are shown in Table S2. Normally distributed data are described with mean and s.d. and analyzed using two-tailed unpaired Student’s t-tests for 2-group comparisons or one-way analysis of variance (ANOVA) with Turkey’s test for multiple-group comparisons. When data deviate from normal distributions (i.e., Fig. 4D), medians with the interquartile range in parentheses are used; statistical analyses are performed with the nonparametric Kruskal-Wallis test for multiple comparisons. Data in Fig.4B is analyzed by Fisher’s exact test. *P* < 0.05 is considered statistically significant.

## List of Supplementary Materials

Materials and Methods

Fig. S1 to S11

Table S1 to S2

## Acknowledgments

The authors acknowledge the use of facilities and instrumentation supported by NSF through the Cornell University Materials Research (CCMR) Science and Engineering Center DMR-1719875; the authors acknowledge Ellis R. Loew for help with ABR. We thank Drs. Stephen I. Pelton and Vishakha Sabharwal at Boston University Chobanian & Avedisian School of Medicine and Boston Medical Center for gifting us the clinical isolate of non-typeable Haemophilus influenzae and Streptococcus pneumoniae.

## Funding

National Institute on Deafness and Other Communication Disorders grant NIHDC016644 (PV, GS)

## Author contributions

Conceptualization: RY, WT

Methodology: WT, XM, CMM, SSL, PC, HS

Investigation: RY, WT, KP

Visualization: WT

Funding acquisition: RY

Supervision: RY

Writing – original draft: RY, WT

Writing – review & editing: RY, WT, CMM

## Competing interests

The authors declare no competing interests.

## Data and materials availability

The main data supporting the results of this study are available within the paper and its Supplementary Information. The raw data generated in this study are available from the corresponding author upon reasonable request.

## Notes

### Competing Interest Statement

The authors have declared no competing interest.

