## Supplementary material for "Harnessing Neutrophils to Deliver Antibiotics Non-Invasively across the Tympanic Membrane for Otitis Media Treatment": Fig. S1-S11 and Table S1-S2

**List of Supplementary Materials**

Materials and Methods

Fig. S1. Neutrophils in the tympanic membrane and middle ear fluid of chinchillas with acute otitis media.

Fig. S2. C3b fragments in the MEF and serum of animals with AOM, and their identification in the protein corona of DSPG/sLip.

Fig. S3. Size and zeta potential distributions of liposomes.

Fig. S4. Gating strategy for neutrophil subsets for the analysis of ex vivo and in vivo liposomes uptake.

Fig. S5. The estimated fluorescence intensity of liposomes on TM and in the neutrophils from MEF.

Fig. S6. Fluorescent images of TM with AOM, treated with DSPG/sLip/DiI.

Fig. S7. Additional results of ex vivo permeation of ciprofloxacin across the TM and in vivo efficacy.

Fig. S8. Rheology of the liposome-containing hydrogel formulations.

Fig. S9. Cell cytotoxicity of the formulations.

Fig. S10. Impact on the bony walls of chinchillas by the treatment of DSPG/sLip/Cip-P407.

Fig. S11. Impact on the hearing sensitivity of healthy chinchillas by the treatment of DSPG/sLip/Cip-P407.

Table S1. Size and zeta potential of liposomes.

Table S2. Normality test of data with Shapiro-Wilk Test.

References (*96, 97*) (numbers for references only cited in SM)

### SUPPLEMENTARY MATERIALS AND METHODS

#### Materials

1,2-distearoyl-sn-glycero-3-phospho-(1'-rac-glycerol) (sodium salt) (DSPG), 1,2-dioleoyl-3-trimethylammonium-propane (chloride salt) (DOTAP), cholesterol, and 1,2-distearoyl-sn-glycero-3-phosphoethanolamine-N-[methoxy(polyethylene glycol)-2000] (ammonium salt) (mPEG$_{2000}$-DSPE, 880120) were obtained from Avanti Polar Lipids Co. Ltd (Birmingham, AL); sephadex® G-50, ciprofloxacin hydrochloride, 1,1′-dioctadecyl-3,3,3′,3′-tetramethylindocarbocyanine perchlorate (41085-99-8), cobra venom factor (Naja naja kaouthia), anti-neutrophil elastase antibody, and poly(ethylene glycol)-block-poly(propylene glycol)-block-poly(ethylene glycol) (Poloxamer 407) were purchased from Sigma Co. Ltd (St. Louis, MO); complement C3 polyclonal antibody (PA1-29715), rabbit anti-goat IgG (H+L) secondary antibody and HRP were from ThermoFisher Co. Ltd (Waltham, MA); 4–20% precast polyacrylamide gel, 2X laemmli sample buffer and precision plus protein dual color standards (#1610374) were from Bio-Rad Laboratories, Inc. (Hercules, CA). CCK-8 Cell Counting Kit (DBOC00128) was bought from VitaScientific Co. Ltd (US); LIVE/DEAD™ Viability/Cytotoxicity Kit (L3224) was from ThermoFisher Co. Ltd (US).

#### Preparation of liposomes

Liposomes were prepared through thin film hydration and extrusion method as previously described *(96)*. The negatively charged liposomes composed of DSPG (52% molar ratio), cholesterol (43% molar ratio), and mPEG$_{2000}$-DSPE (5% molar ratio) were dissolved in a mixed solvent (chloroform/ethanol/deionized water) and distilled under vacuum until a thin film was formed. The film was hydrated with 0.32 M ammonium sulfate at 60°C and extruded through 400-nm, 200-nm, and 100-nm polycarbonate membranes sequentially, 8 to 10 times for each membrane. Ciprofloxacin was loaded into the liposomes by the established ammonium sulfate gradient method at 60ºC (in a water bath) *(97)*. Positively charged liposomes composed of 32% DOTAP, 20% HSPC, 43% cholesterol, and 5% mPEG$_{2000}$-DSPE were made following a similar method as described above, except the dissolution step that was performed in chloroform instead of a mixture solvent. DiI dye (100 μg/ml) was added to the lipids before vacuum distillation. The size and zeta potential of liposomes (diluted 20 times from the aforementioned solution with deionized water) were analyzed with Dynamic Light Scattering (DLS) (Nano ZS90, Malvern, UK); The amount of ciprofloxacin in liposomes was quantified by High-Performance Liquid Chromatography (HPLC, Shimadzu).

#### Auditory brainstem response (ABR) measurements

ABR experiments were conducted with a custom-designed stimulus generation and measurement system built around National Instruments software (LabVIEW) and hardware. The hardware included a GPIB controller and an ADC board. A custom LabVIEW program computed the stimuli and downloaded the stimuli to a programmable stimulus generator (Hewlett Packard 33120A). The stimulus was then filtered by an antialiasing filter (KrohnHite 3901) and attenuated (Tucker-Davis Technologies). The filter and the attenuator were controlled by the LabView software. Simultaneous with stimulus output, the 2 ADC channels sampled the amplified ABR signal and the output of a microphone sealed in the ear canal of the animal.
The attenuated stimulus was played through a hearing-aid earphone placed within the intact ear canal of adult female chinchillas (400-600 g) anesthetized by inhalation of isoflurane (60 mg per

kg). The earphone coupler included a microphone that monitored the sound stimulus level. ABR signals, obtained in a sound-attenuating booth, were measured with a differential amplifier with a gain of 10,000 and a measurement bandwidth of 100 Hz to 3 kHz. The measurements were obtained from the positive electrode in the muscle behind the measured ear; the negative electrode was at the cranial vertex, and the ground electrode behind the contralateral ear. After obtaining baseline measurements, 200 µl of DSPG/sLip/Cip-P407[15%] gel was applied to the ear canal, and the same measurements were immediately performed to examine the resultant changes, if any, in auditory sensitivity thresholds.

**NTHi/*S. Pneumoniae* coinfection OM model**

All procedures and manipulations were performed in accordance with the approved protocol by the IACUC and the guidelines by Cornell University CARE. Isolates of NTHi and *Streptococcus Pneumoniae* grown to the mid-log phase were diluted in Hanks balanced salt solution (HBSS), and ~100 to 200 CFU in 100 µl for each pathogen were introduced directly into each middle ear bullae under aseptic conditions. Daily otomicroscopy was performed to monitor the bulging TM. Once abnormality was identified, the middle ear cavity was accessed 48 to 72 hours later. TMs of the animals to receive the formulation were observed with the speculum of an otoscope, after which the liquid hydrogel was injected through the speculum using a soft 18-gauge, 1.75-inch catheter. The experiment was terminated on day 1 due to the development of pneumonia in the infected animals. The amount of Cip in the middle ear was quantified with HPLC.

**Histopathology**

Formulations were administered to the ear canals of live NTHi-infected OM chinchillas. Seven days later, they were euthanized. After sacrifice, the TMs were excised and immediately fixed with 10% neutral buffered formalin overnight, then decalcified, embedded in paraffin, sectioned (5 µm thick), and stained with Hematoxylin and Eosin (H&E) by the Section Anatomic Pathology Histology Laboratory at Cornell University (Ithaca, New York, USA). All stained specimens were evaluated under Keyence Box Microscope (BZ-180, Itasca, IL, US), and the thickness of TMs was measured by Image J software (Bethesda, Maryland, US).

### SUPPLEMENTARY FIGURES

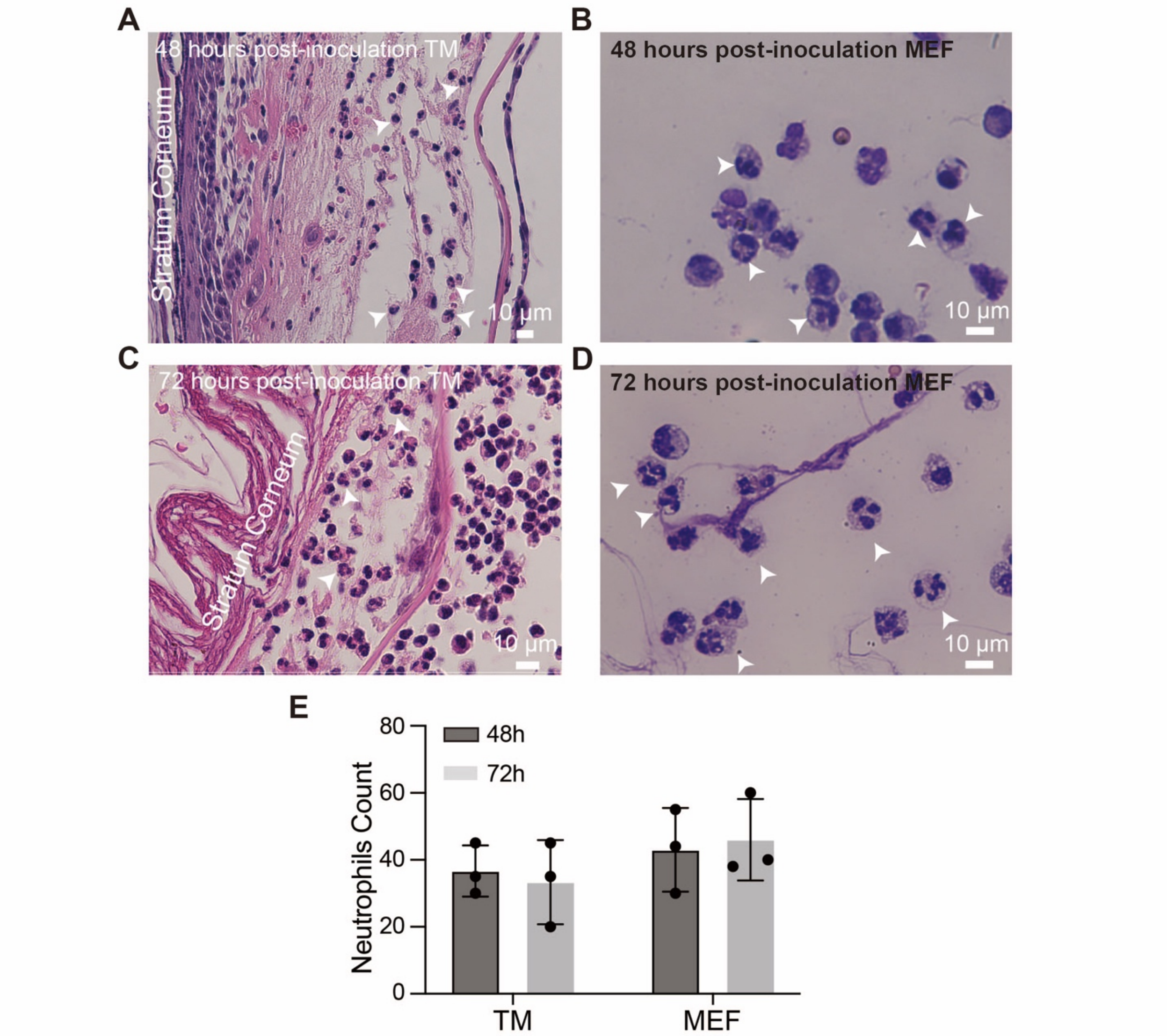


**Fig. S1. Neutrophils in the tympanic membrane and middle ear fluid of chinchillas with acute otitis media.** (**A-B**) Hematoxylin and eosin (H&E)-stained sections of the tympanic membrane (TM) (A) and Wright-Giemsa-stained middle ear fluid (MEF) (B) excised from chinchillas with acute otitis media (AOM), i.e., two days after the NTHi inoculation. The white arrows indicate the neutrophils residing within the stratum corneum layer and the connective tissue and the mucosal epithelium in (A) and middle ear fluid in (B). (**C-D**) H&E-stained TM (C) and Wright-Giemsa-stained MEF (D) sampled from the animals after three days of NTHi inoculation. The white arrows indicate the neutrophils residing within the stratum corneum layer in (C) and middle ear fluid in (D). (**E**) The statistical analysis of the neutrophil levels in the tympanic membrane and middle ear fluid at 48 h and 72 h post-inoculation, based on three histological images (of a 280 μm × 370 μm area) for each group. Scale bar, 10 μm.

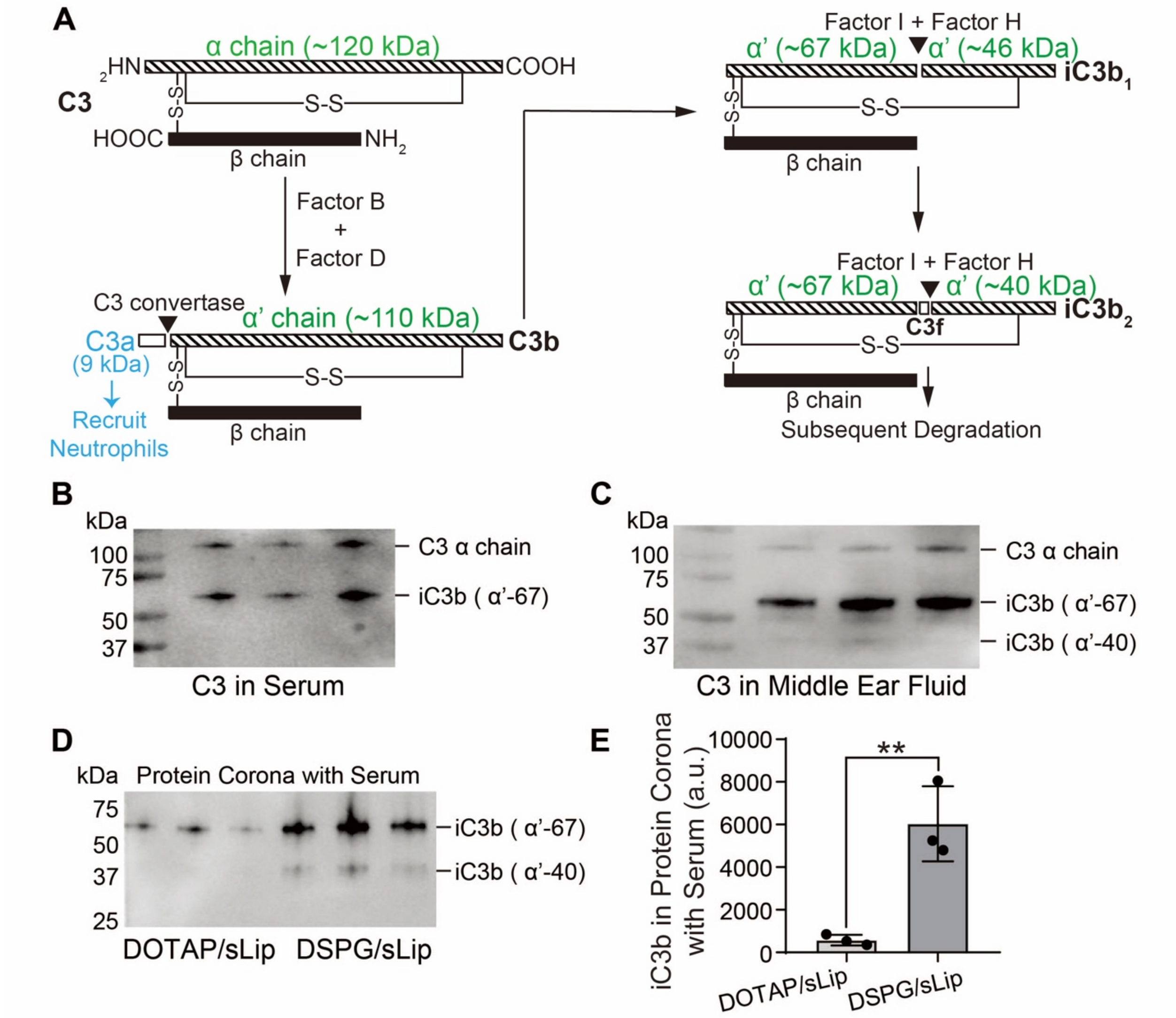


**Fig. S2. C3b fragments in the MEF and serum of animals with AOM, and their identification in the protein corona of DSPG/sLip.** (**A**) Schematic of the alternative activation pathway that leads to C3 cleavage. C3 comprises two polypeptide chains (α and β) linked by disulfide bonds. The molecular weights of C3 α chain, C3a (that recruits neutrophils), and C3b α' chain are ~120 kDa, ~9 kDa, and ~110 kDa, respectively. However, the C3b α'fragment is rarely captured using western blot because the C3b fragment is often cleaved by Factor I and Factor H into iC3b, with molecular weights of ~67 kDa for the iC3b α'-67 chain, ~46 kDa for the iC3b α'-46 chain, and ~40 kDa for the iC3b α'-40 chain, respectively. (**B-C**) Western blot of the C3b fragments in the serum (B) and MEF (C) of chinchillas with AOM. (**D-E**) Western blot of the liposome protein coronas formed by incubating DOTAP/sLip or DSPG/sLip with the serum collected from animals with AOM and the quantity of iC3b fragments in their protein coronas, respectively. $^{**}P = 0.0082$ by two-tailed unpaired Student's t-test. Data are presented as mean ± SD (n = 3).

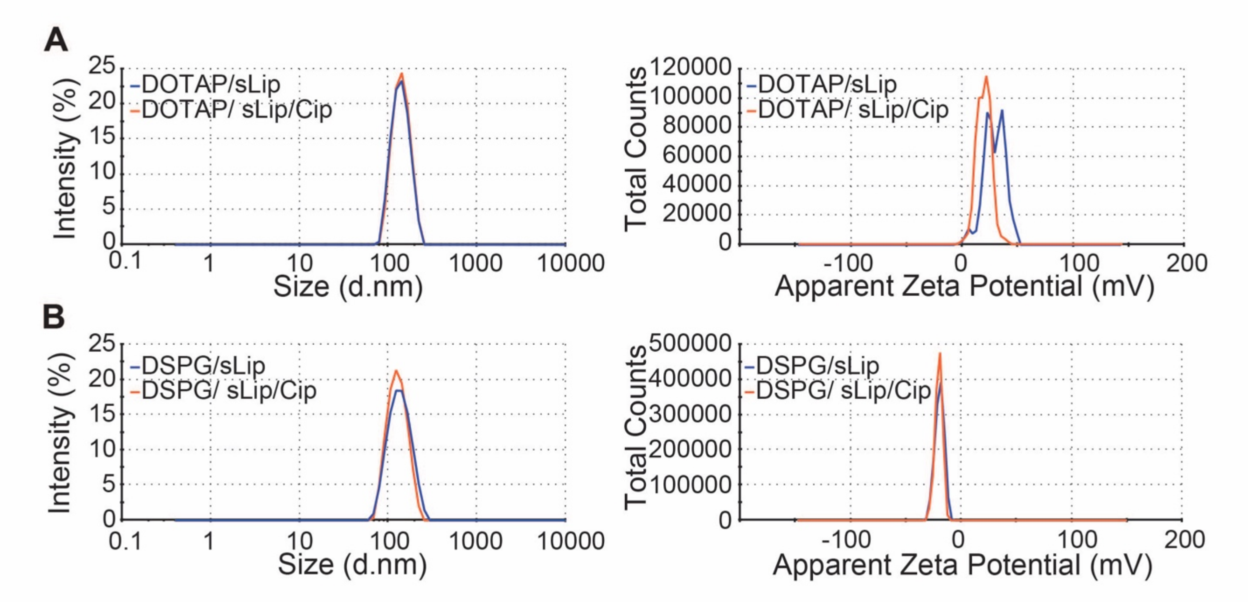


**Fig. S3. Size and zeta potential distributions of liposomes.** (**A**) Size and zeta potential distributions of DOTAP/sLip and DOTAP/sLip/Cip. (**B**) Size and zeta potential distributions of DSPG/sLip and DSPG/sLip/Cip.

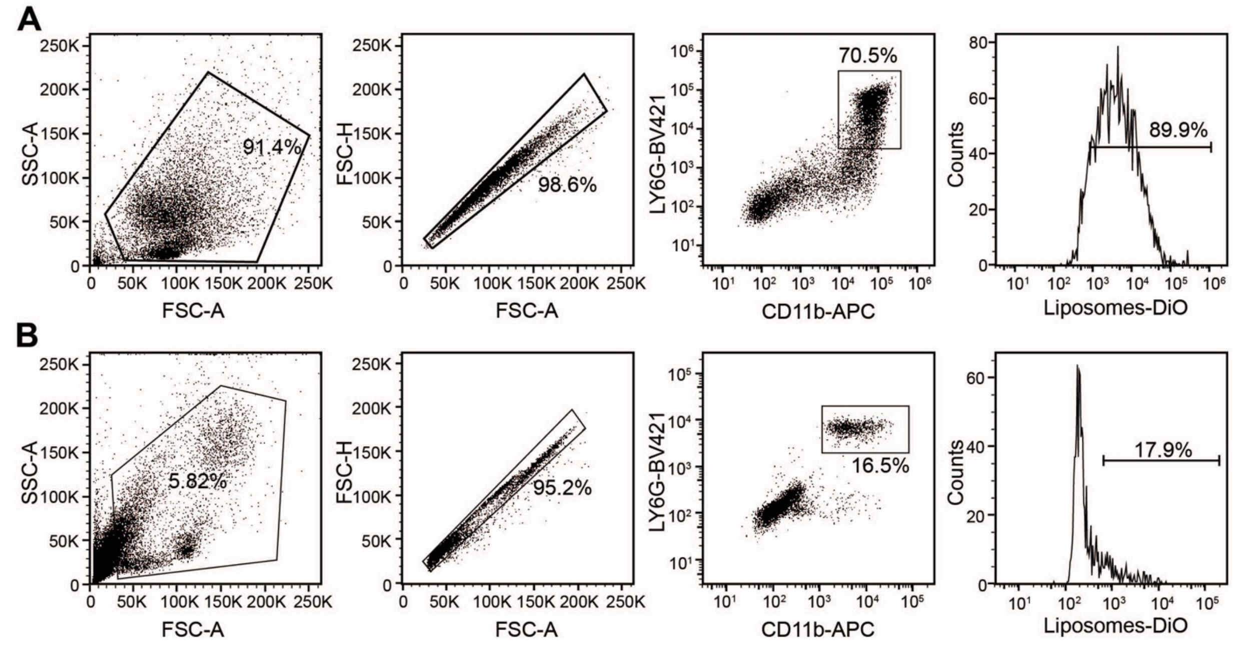


**Fig. S4. Gating strategy for neutrophil subsets for the analysis of ex vivo and in vivo liposomes uptake.** (**A**) Gating strategy for flowcytometric analysis of neutrophils internalization of liposomes ex vivo in Fig. 1C. (**B**) Gating strategy for flowcytometric analysis of neutrophils internalization of liposomes in vivo in Fig. 1D.

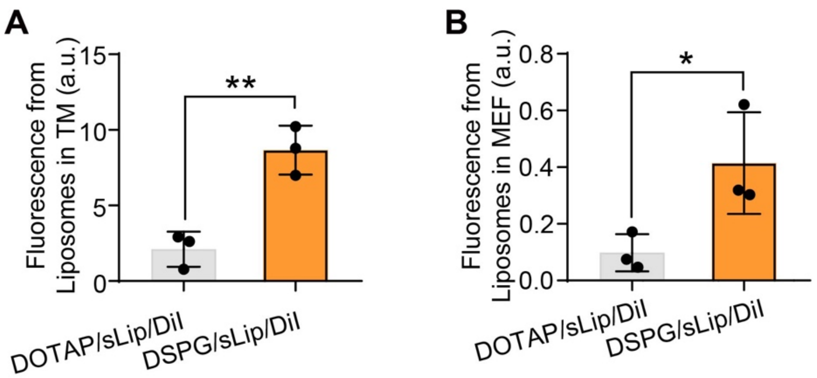


**Fig. S5. The estimated fluorescence intensity of liposomes on TM and in the neutrophils from MEF.** (**A**) The fluorescence intensity of DOTAP/sLip/DiI and DSPG/sLip/DiI on TM calculated from Fig. 1E. (**B**) The fluorescence intensity of DOTAP/sLip/DiI and DSPG/sLip/DiI in neutrophils from MEF calculated from Fig. 1F. Data are presented as mean ± SD (n = 3). *P* values were determined by two-tailed unpaired Student's t-tests; $*P < 0.05$; $**P < 0.01$.

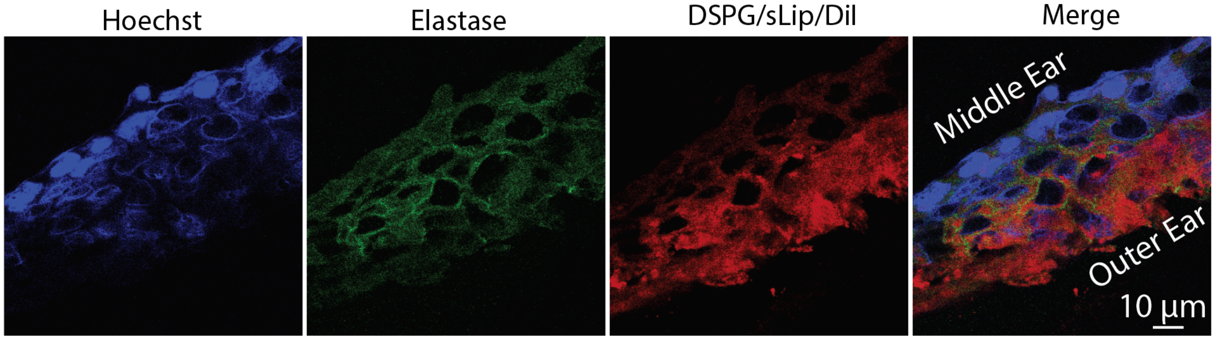


**Fig. S6. Fluorescent images of TM with AOM, treated with DSPG/sLip/DiI.** Fluorescent images of the tympanic membrane of animals with acute otitis media treated with DSPG/sLip/DiI, where the neutrophil elastase is stained in green, showing its penetration into the tympanic membrane. Scale bar, 10 µm.

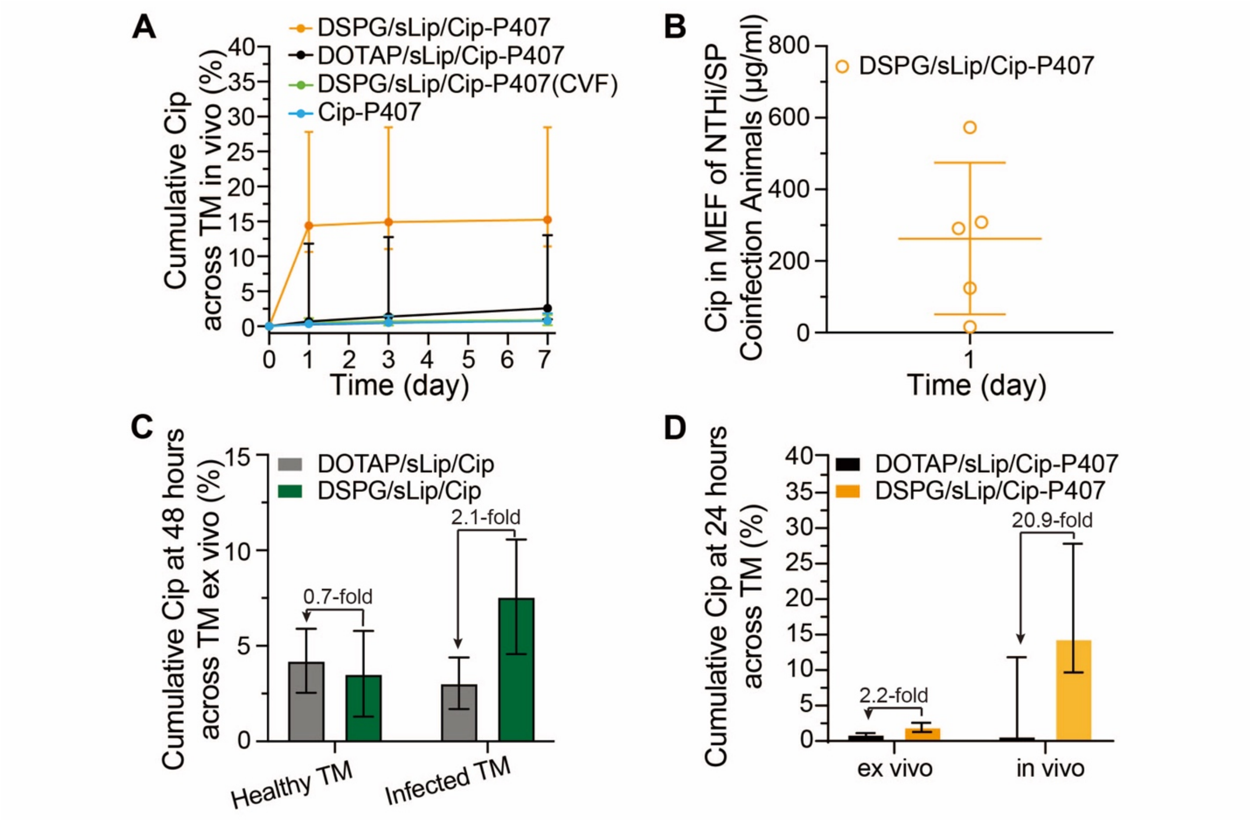


**Fig. S7. Additional results of ex vivo permeation of ciprofloxacin across the TM and in vivo efficacy.** (**A**) Percentage of the ciprofloxacin (Cip) contained in the hydrogel formulation that permeated across the TM over the course of the 7-day treatment (n = 4). Data are medians with the interquartile ranges. (**B**) Concentration of Cip in the MEF of animals infected with NTHi and *Streptococcus pneumoniae* simultaneously and treated with DSPG/sLip/Cip-P407 for 24 hours (n = 5). Data are presented as mean ± SD. (**C**) Permeation (%) of Cip across the TMs of healthy chinchillas (n = 3) and chinchillas with AOM (n = 4) after 48 hours of treatment using DSPG/sLip/Cip and DOTAP/sLip/Cip. Data are presented as mean ± SD. (**D**) Permeation (%) of Cip across the TM of animals with AOM after 24 hours of treatment using DSPG/sLip/Cip-P407 and DOTAP/sLip/Cip-P407, measured using excised auditory bullae, i.e., ex vivo (n = 4), or by extracting the MEF of infected and treated animals, i.e., in vivo (n = 4). Data are presented as medians with the interquartile ranges.

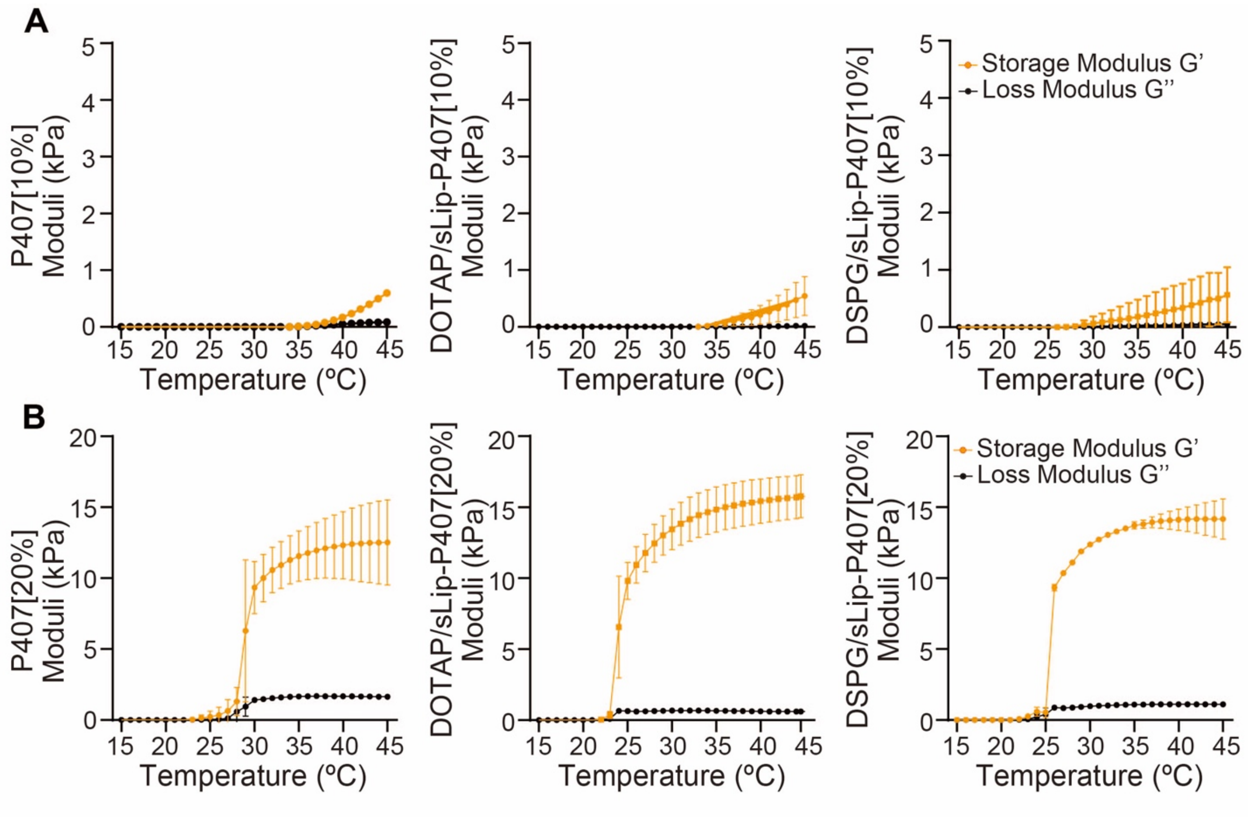


**Fig. S8. Rheology of the liposome-containing hydrogel formulations.** (**A**) Rheology of P407[10%], DOTAP/sLip-P407[10%] and DSPG/sLip-P407[10%], as a function of temperature. All formulations contain 10% (w/v) P407. (**B**) Rheology of P407[20%], DOTAP/sLip-P407[20%] and DSPG/sLip-P407[20%], as a function of temperature. All formulations contain 20% (w/v) P407. Data are presented as mean ± SD (n = 3).

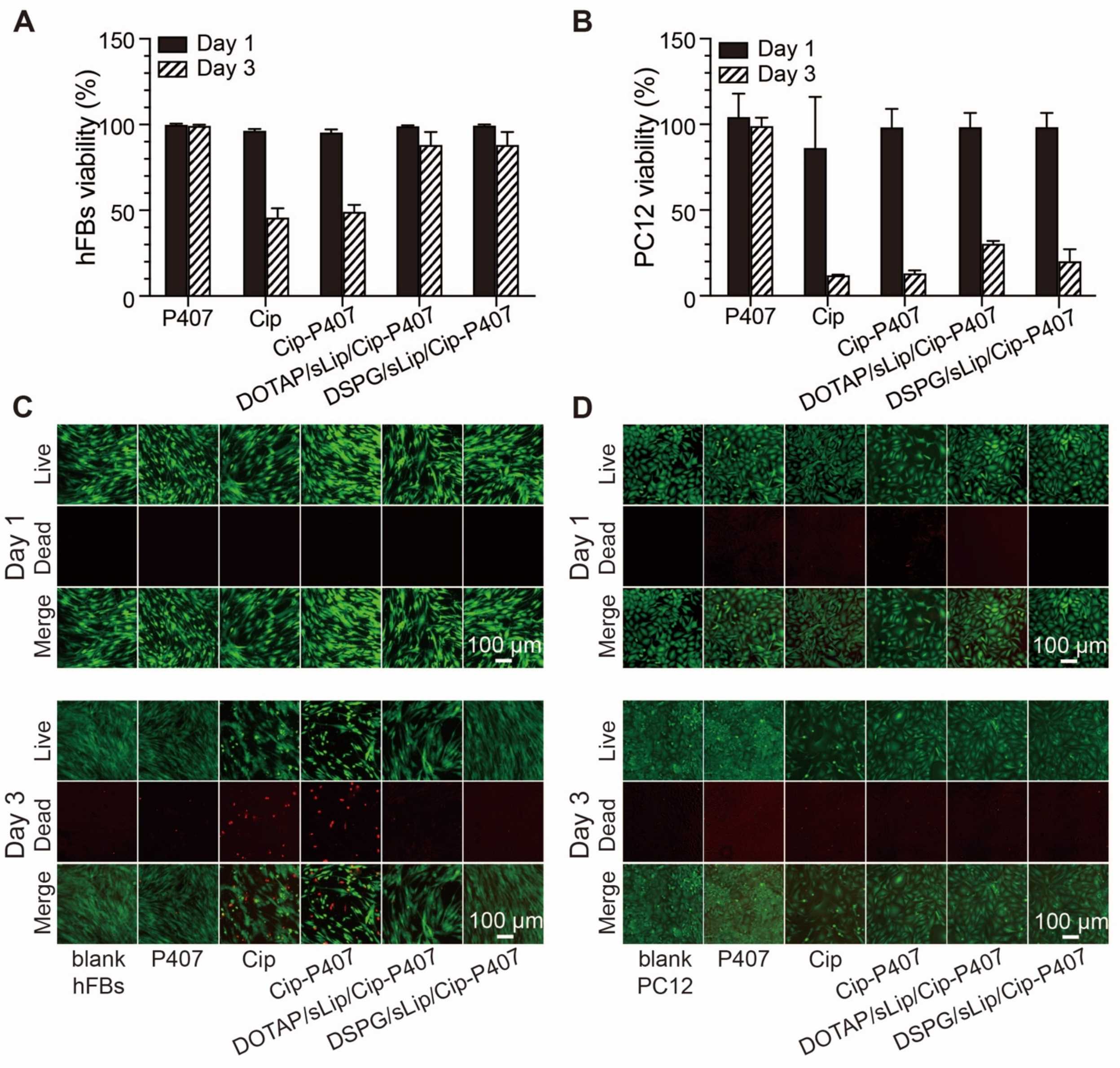


**Fig. S9. Cell cytotoxicity of the formulations**. (**A-B**) Survival rates (determined by MTS assay) of human dermal fibroblasts (hFBs) (A) and PC12 cells (B) after incubating with Cip, P407, Cip-P407, DOTAP/sLip/Cip-P407, and DSPG/sLip/Cip-P407. Data are presented as mean ± SD (n = 3). (**C-D**) LIVE/DEAD assay of hFBs (C) and PC12 (D), which was done to confirm the data in (A) and (B), after incubation for 1 or 3 days with Cip, P407, Cip-P407, DOTAP/sLip/Cip-P407, and DSPG/sLip/Cip-P407, respectively. GFP (green fluorescent protein), live cells (green); TRITC (tetramethyl rhodamine isothiocyanate), dead cells (red). Scale bar, 100 μm. All Cip-containing formulations contain 0.5 mg of Cip; all P407-containing formulations contain 15% (w/v) P407.

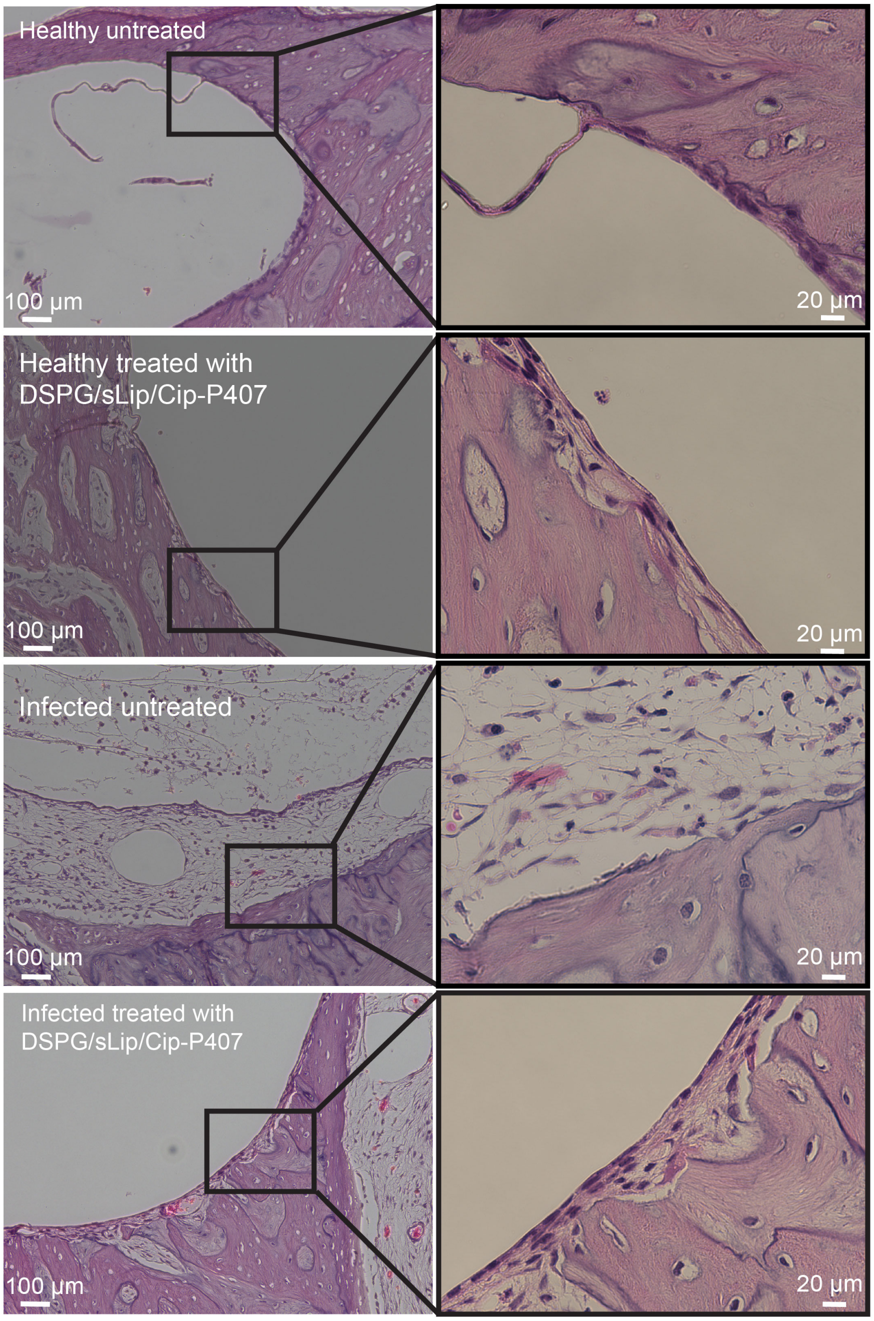


**Fig. S10. Impact on the bony walls of chinchillas by the treatment of DSPG/sLip/Cip-P407.** Auditory bullae harvested from chinchillas that are healthy untreated, healthy treated with DSPG/sLip/Cip-P407 for 7 days, infected untreated, and infected and treated with DSPG/sLip/Cip-P407 for 7 days. In all formulations containing ciprofloxacin, its amount is 0.5 mg Cip; in all formulations containing P407, its concentration is 15% (w/v). Scale bar, 100 µm and 20 µm.

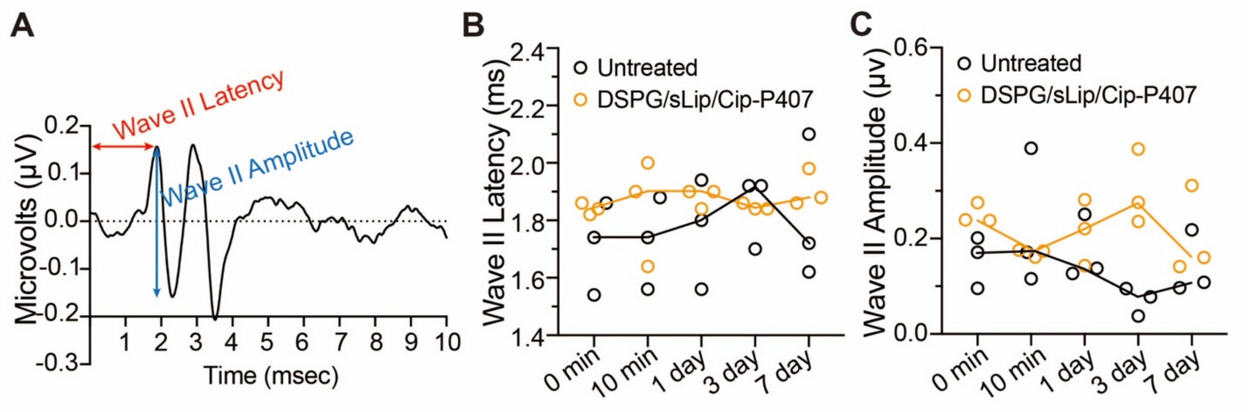


**Fig. S11. Impact on the hearing sensitivity of healthy chinchillas by the treatment of DSPG/sLip/Cip-P407.** (**A**) Auditory brainstem response (ABR) waves were recorded over 10 ms in response to an acoustic click. The red and blue arrows indicate wave II absolute latency and amplitude, respectively. (**B-C**) Variation of wave II latency (B) and amplitude (C) over the course of 7 days in untreated animals and ones treated with DSPG/sLip/Cip-P407 (n = 3). The black and orange circles are individual data points; solid lines indicate the mean values. The concentration of P407 is 15% (w/v).

### SUPPLEMENTARY TABLES

**Table S1. Size and zeta potential of liposomes**. Liposomes, 20 times diluted with deionized water, were tested with DLS for size (nm) and surface zeta potential (mv). Data are presented as mean ± SD (n = 3).

| **Liposomes** | **Size (nm)** | **Zeta Potential (mv)** |
|---|---|---|
| DOTAP/sLip | 134.2 ± 1.5 | +26.8 ± 1.5 |
| DOTAP/sLip/DiI | 131.1 ± 5.5 | +23.7 ± 3.0 |
| DOTAP/sLip/Cip | 130.8 ± 5.2 | +20.1 ± 0.6 |
| DOTAP/sLip/DiI/Cip | 132.5 ± 4.3 | +23.2 ± 4.0 |
| DSPG/sLip | 127.4 ± 5.2 | –20.1 ± 0.4 |
| DSPG/sLip/DiI | 132.5 ± 0.7 | –20.8 ± 0.7 |
| DSPG/sLip/Cip | 135.8 ± 0.7 | –20.7 ±1.5 |
| DSPG/sLip/DiI/Cip | 130.7 ± 4.7 | –21.0 ± 1.1 |

**Table S2. Normality test of data with Shapiro-Wilk Test.** The Shapiro–Wilk test is used to test the normality of the data. $P > 0.05$, normal distribution; $P < 0.05$, asymmetrical distribution.

| **Data** | ***P* Value (Shapiro-Wilk Test)** |
|---|---|
| *Fig2b* | |
| DOTAP/sLip/Cip | 1 |
| DSPG/sLip/Cip | 0.9991 |
| Cip | 0.2974 |
| Cip-P407 | 0.1612 |
| *Fig2c* | |
| DSPG/sLip/Cip | 0.7441 |
| DSPG/sLip/Cip-P407 | 0.7876 |
| DOTAP/sLip/Cip | 0.879 |
| DSPG/sLip-Cip-P407 | 0.2706 |
| DOTAP/sLip/Cip-P407 | 0.9999 |
| Cip-P407 | 0.1494 |
| *Fig4c* | |
| Untreated | 0.3345 |
| Cip-P407 | 0.3429 |
| DOTAP/sLip/Cip-P407 | 0.2254 |
| DSPG/sLip-P407 | 0.4097 |
| DSPG/sLip/Cip-P407 | [#]0 |
| *Fig4d* | |
| DSPG/sLip/Cip-P407 | 0.3205 |
| DOTAP/sLip/Cip-P407 | [*]0.02688 |
| Cip-P407 | 0.7955 |
| *Fig5d* | |
| DSPG/sLip/Cip-P407(CVF) | 0.3177 |
| *Fig5e* | |
| DSPG/sLip/Cip-P407(CVF) | 0.2131 |

[#]Data are 0, 0, 0, and 0

[*]Asymmetrical
